# The evolutionary history of *Drosophila simulans* Y chromosomes reveals molecular signatures of resistance to sex ratio meiotic drive

**DOI:** 10.1101/2022.12.22.521550

**Authors:** C Courret, D Ogereau, C Gilbert, A.M Larracuente, C Montchamp-Moreau

## Abstract

The recent evolutionary history of the Y chromosome in *Drosophila simulans*, a worldwide species of Afrotropical origin, is closely linked to that of X-linked meiotic drivers (Paris system). The spread of the Paris drivers in natural populations has elicited the selection of drive resistant Y chromosomes. To infer the evolutionary history of the Y chromosome in relation to the Paris drive, we sequenced 21 iso-Y lines, each carrying a Y chromosome from a different location. Among them, 13 lines carry a Y chromosome that is able to counteract the effect of the drivers. Despite their very different geographical origins, all sensitive Y’s are highly similar, suggesting that they share a recent common ancestor. The resistant Y chromosomes are more divergent and segregate in four distinct clusters. The phylogeny of the Y chromosome confirms that the resistant lineage predates the emergence of Paris drive. The ancestry of the resistant lineage is further supported by the examination of Y-linked sequences in the sister species of *D. simulans, D. sechellia,* and *D. mauritiana*. We also characterized the variation in repeat content among Y chromosomes and identified multiple simple satellites associated with resistance. Altogether, the molecular polymorphism allows us to infer the demographic and evolutionary history of the Y chromosome and provides new insights on the genetic basis of resistance.

## INTRODUCTION

In *Drosophila melanogaster* and *D. simulans*, like in most Drosophila species, the Y chromosome is entirely heterochromatic, composed of transposable elements and other repetitive sequences such as satellite DNA (Bachtrog 2013). In both species the Y chromosome represents 20% of the male genome (around 40 Mb) (Hoskins et al. 2002) but contains only a handful of genes, expressed during spermatogenesis and necessary for male fertility (Kennison 1981; Bernardo Carvalho, Koerich, et Clark 2009). Indeed, XO males are usually viable but sterile. Beyond its essential role in male fertility, the Y chromosome of *D. melanogaster* has been shown to impact male fitness through a variety of phenotypic effects, on behavior or thermotolerance (Huttunen et Aspi, 2003.; Rohmer et al. 2004). In fact, variation in Y-linked heterochromatin appears able to modulate gene expression across the whole genome by modifying the chromatin landscape (Brown, Nguyen, et Bachtrog 2020; Alan T. Branco, Brito, et Lemos 2017; Lemos et al. 2014; Francisco et Lemos 2014). Similar effects are likely in *D. simulans* (Branco et al. 2013).

The large phenotypic variation caused by the Y chromosome of *D. melanogaster* is in sharp contrast with its very low nucleotide diversity in gene sequence compared to other chromosomes (Zurovcova et Eanes 1999; Larracuente et Clark 2013). In *D. simulans,* the sibling species of *D. melanogaster*, the gene nucleotide diversity of the Y chromosome is even lower (Zurovcova et Eanes 1999; Kopp, Frank, et Fu 2006; Helleu et al. 2019).

This low variation can be explained by multiple factors. First, the Y chromosome has a reduced effective population size compared to the rest of the genome (one quarter of that of the autosomes). Second, because of its hemizygous state, the fitness conditions allowing the maintenance of stable, non-neutral polymorphisms are very restrictive (Clark 1987; Clark 1990). Finally, the Y chromosome does not recombine, which causes natural selection on a Y-linked locus to affect the entire chromosome. It follows that beneficial mutation can lead to the fixation of the carrier Y chromosome in the population (Charlesworth, Coyne, and Barton 1987), while a deleterious mutation can lead to its loss. Population genetics models thus suggest that neutral variations within populations should be rarely maintained whereas inter-population differences can rapidly accumulate (Clark 1987). Thus, the *Drosophila* Y chromosome is expected to be particularly sensitive both to the demographic history of populations and to selection events. Indeed, in *D. melanogaster*, the ancestral African populations show significantly less Y-linked variation than the cosmopolitan ones, which could be explained by the demographic history of the species and recent natural selection in Africa (Larracuente et Clark 2013).

Y chromosome evolution can also be shaped by intragenomic conflict triggered by X-linked meiotic drivers. These favor the transmission of X chromosomes bearing the driver (hereafter X^SR^), thus, increasing the proportion of females in progeny and inducing a deviation from the balanced - also called Fisherian - sex ratio. *Sex Ratio* (*SR*) systems are widespread in the *Drosophila* genus, with 19 *SR* systems known in 16 species, among which 3 systems have been described in *Drosophila simulans* (Courret et al. 2019). During spermatogenesis, an X^SR^ typically prevents the formation of functional Y bearing sperm, resulting in a strong female bias in the progeny of carrier males. Thus, the effective size of the Y chromosome is drastically reduced in populations invaded by X^SR^. Following the spread of X^SR^, natural selection strongly favors any variant that can counteract the effect of the driver and restore a balanced sex ratio (reviewed in Helleu, Gérard, et Montchamp-Moreau 2015). Such drive suppressors are expected to evolve at unlinked loci, *i.e.*, on autosomes and the Y chromosome. Resistant Y chromosomes have direct and strong selective advantage and thus are expected to quickly reach fixation (Clark 1987; Hall 2004).

*Drosophila simulans* is particularly well suited to investigate the impact of SR drive on the evolutionary history of the Y chromosome. In this worldwide species, autosomal suppressors and resistant Y chromosomes have evolved in response to the spread of X-linked drivers known as the Paris system (Cazemajor, Landré, et Montchamp-Moreau 1997). The Paris system likely emerged less than 500 years ago in East Africa (Fouvry et al. 2011; Bastide et al. 2011). It then spread to islands of the Indian Ocean and in Sub-Saharan Africa (Bastide et al. 2013). Drive suppression is complete over its geographic range and the frequency of X^SR^ was found to decrease in some locations, like in Madagascar, Kenya, or Mayotte (Bastide et al. 2011). During the last decade, the invasion of X^SR^ was observed in North African populations, where sensitive Y chromosomes were then replaced by resistant ones within a few years (Helleu et al. 2019). On the whole, in *D. simulans*, the Y chromosomes appeared to be highly polymorphic for their resistance ability with a continuum of phenotypes, from complete resistance to high sensitivity (Montchamp-Moreau, Ginhoux, et Atlan 2001). In contrast, they exhibit a very low level of gene sequence variation, with only three single nucleotide polymorphisms (SNPs) detected along a total of 13 kb of gene fragments; those define three haplotypes - TTA, TTT, CAA - the latter of which is associated with resistance to Paris *SR* drive (Helleu et al. 2019).This low number of SNPs strongly suggests that the phenotypic variation results from variation in repeated sequences and/or from structural variations, both of which are extensive on the Y chromosomes of *D. simulans* (Helleu et al. 2019; Chang et al. 2022). Surprisingly, the CAA haplotype appeared to be ancestral in the *simulans* clade (Helleu et al. 2019), which suggests that resistant Y chromosomes have long predated the emergence of the Paris driver (Helleu et al. 2019). Consistently with this scenario, the Y chromosome of its sibling, *D. sechellia,* is resistant to Paris X^SR^ (Tao et al; 2007).

Thanks to the recent advance in long read sequencing, about one third of the Y chromosomes of *D. melanogaster* and *D. simulans* (13 Mb out of 40), are now assembled (Chang et Larracuente 2019; Chang et al. 2022). Here we take advantage of those assemblies to further investigate the molecular variation of the Y chromosome in *D. simulans* in relation to the evolution of the Paris *SR* system. We sequenced 21 iso-Y lines (each with a Y chromosome originating from a single male) from different locations around the world; 8 from the sensitive lineage TTA and 13 from the resistant lineage CAA (Figure 1A). Despite their very different geographical origins all sensitive Y chromosomes appear to be highly similar. The resistant Y chromosomes are more divergent and segregate in four distinct groups. By reconstructing the phylogeny of the Y chromosome, we confirmed that the resistant lineage predates the emergence of the Paris X^SR^. We also examined the variation in repeat content among Y chromosomes. We observed almost no variations in gene and TE copy number, neither of them showing an association with the resistant phenotype. By contrast, extensive variation was found among simple satellites, and we identified a set of satellites which presence and quantity are associated with resistance to the SR system. Altogether this new approach allowed us to better understand the evolutionary history of the Y chromosome in relation with *SR* systems and also yielded new insights on the genetic basis of the resistance.

**Figure 1.**
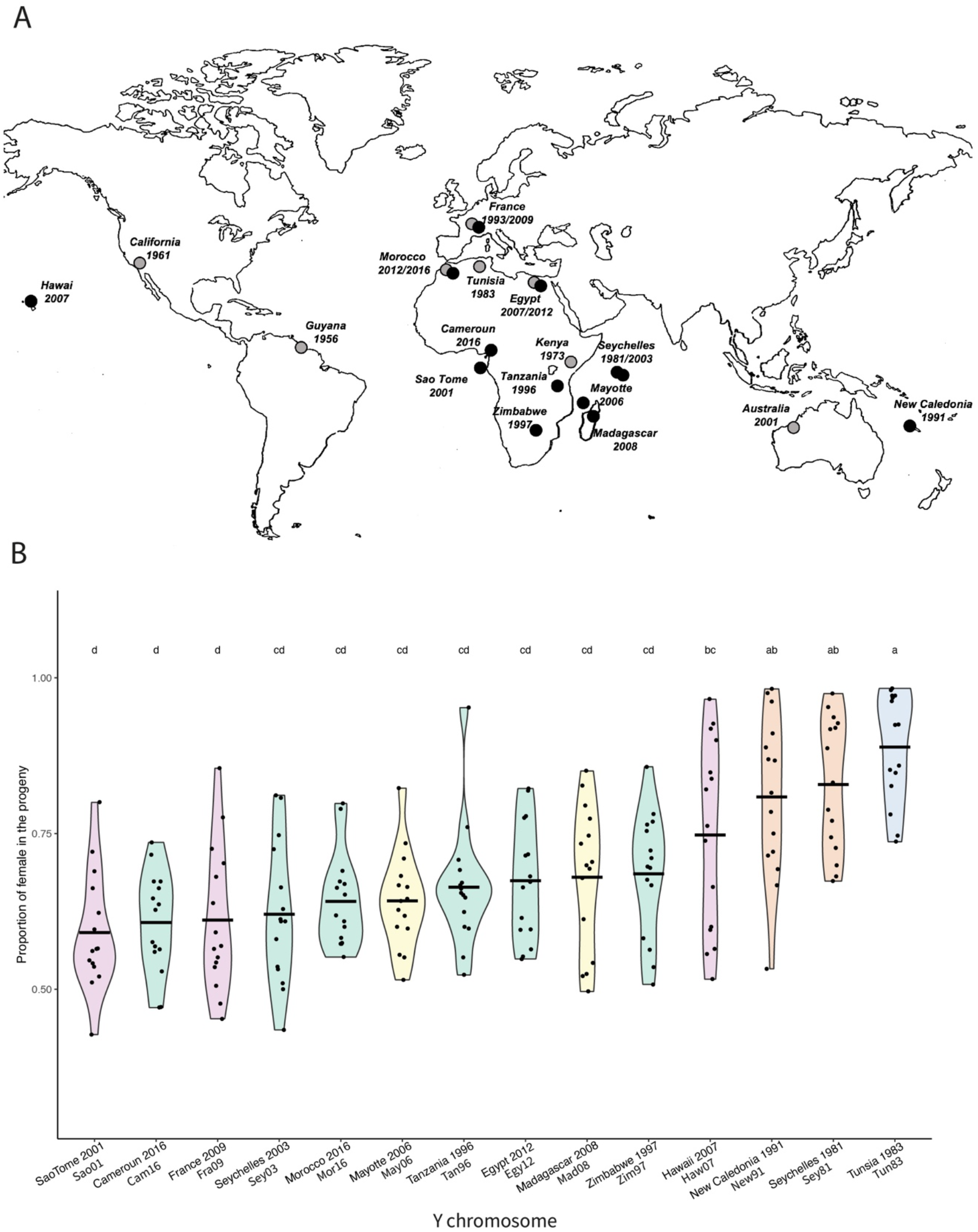
A. Geographical location and date of collection of the Y chromosomes studied (●) Lines carrying a Y chromosome with the CAA haplotype (resistant). (●) Lines carrying a Y chromosome with the TTA haplotype (sensitive). B. Test of the resistance ability of the Y chromosomes. Each violin plot represents the proportion of females in the progeny of males carrying the reference X^SR^ and the Y chromosome from each iso-Y line. Tun83 corresponds to our reference sensitive Y chromosome. Each dot corresponds to the progeny of an individual male. The letters above each violin plot correspond to the group determined by the Tukey HSD post hoc-test performed on an Anova model. Violin plots are colored according to the clusters identified by PCA (same color code as in Figure 2).

## RESULTS

### 1. Resistance ability of the Y chromosomes from the resistant CAA lineage

Among the 21 *D. simulans* Y chromosomes included in the present study, 13 belong to the resistant CAA lineage and were checked for their resistance ability. To do so we produced males carrying the Y chromosome to be tested, a reference X^SR^ chromosome, and suppressor-free autosomes from the reference ST8 line (crossing scheme in Montchamp-Moreau, Ginhoux, and Atlan 2001). For each Y, the individual progeny of 15 males was examined (Figure 1B). We observed significant variation in the resistance ability among the CAA Y chromosomes (ANOVA test, F=12.36, df=13, p<2e-16). The mean sex-ratio in the progeny of carrier males ranged from 60% to 83% depending on the Y chromosome, compared to 89% for the sensitive Y^ST8^ (Figure 1B, Supplementary Table1). Except for Sey81 (Seychelles 1981) and New91 (New Caledonia 1991), which exhibit a low resistance ability with a mean sex-ratio of 81% and 83%, respectively, all CAA Y chromosomes appear significantly more resistant compared to Y^ST8^, our reference sensitive Y chromosome (Figure 1B, Supplementary Table1).

### 2. The molecular variation essentially occurs among resistant Y chromosomes

Using the Y chromosome from the XD1 strain (Chang et al. 2022) as a reference, we identified 8190 Y-linked polymorphic sites among the 21 iso-Y lines sequenced. The global nucleotide diversity is very low (1.808e-04) and most of this diversity comes from the CAA resistant Y chromosomes (Supplementary Figure 1). On average, these share the same allele at 76.89% of the polymorphic sites, while on average the sensitive TTA Y chromosomes share the same alleles at 97.77 % of the detected polymorphic sites (Table1). Sensitive Y chromosomes are thus highly similar despite originating from geographical regions far away from each other. They all mainly share the same allele as the reference allele (Supplementary Figure 1). It follows that the Y chromosome of the XD1 strain (from Winters, California, 1995) belongs to the sensitive TTA lineage.

To examine the population structure of the Y chromosomes, we performed a PCA analysis based on the 8190 SNPs identified. It revealed 5 different clusters. Unsurprisingly, based on their high percentage of identity, all the sensitive Y chromosomes cluster together (Cluster I, Figure 2). The resistant Y chromosomes segregate in 4 different clusters. Cluster II is composed of Egy12, Tan96, Zim97, Mor16, Cam16 and Sey03, all originating from continental Africa or nearby islands. Cluster III is composed of Sey81 and New91, from Seychelles and New Caledonia, respectively. Cluster IV consists of May09 and Mad98, respectively sampled in Mayotte and Madagascar, the area from which the *D. simulans* species is thought to have originated (Lachaise et Silvain, 2004; Dean et Ballard 2004). Cluster V is composed of Fra09, Haw07 and Sao01, which are from very far apart geographical areas (France, Hawaii and SaoTome, respectively). The first axis of the PCA (29.8%) separates Cluster III from all the other Y chromosomes (Figure 2). This means that the divergence between Sey81/New91 and all the other Ys is higher than between all the other resistant Y chromosomes and the sensitive Y chromosomes. Interestingly, those Y chromosomes from Seychelles and New Caledonia are also very similar in resistance ability (Figure 1B).

**Figure 2.**
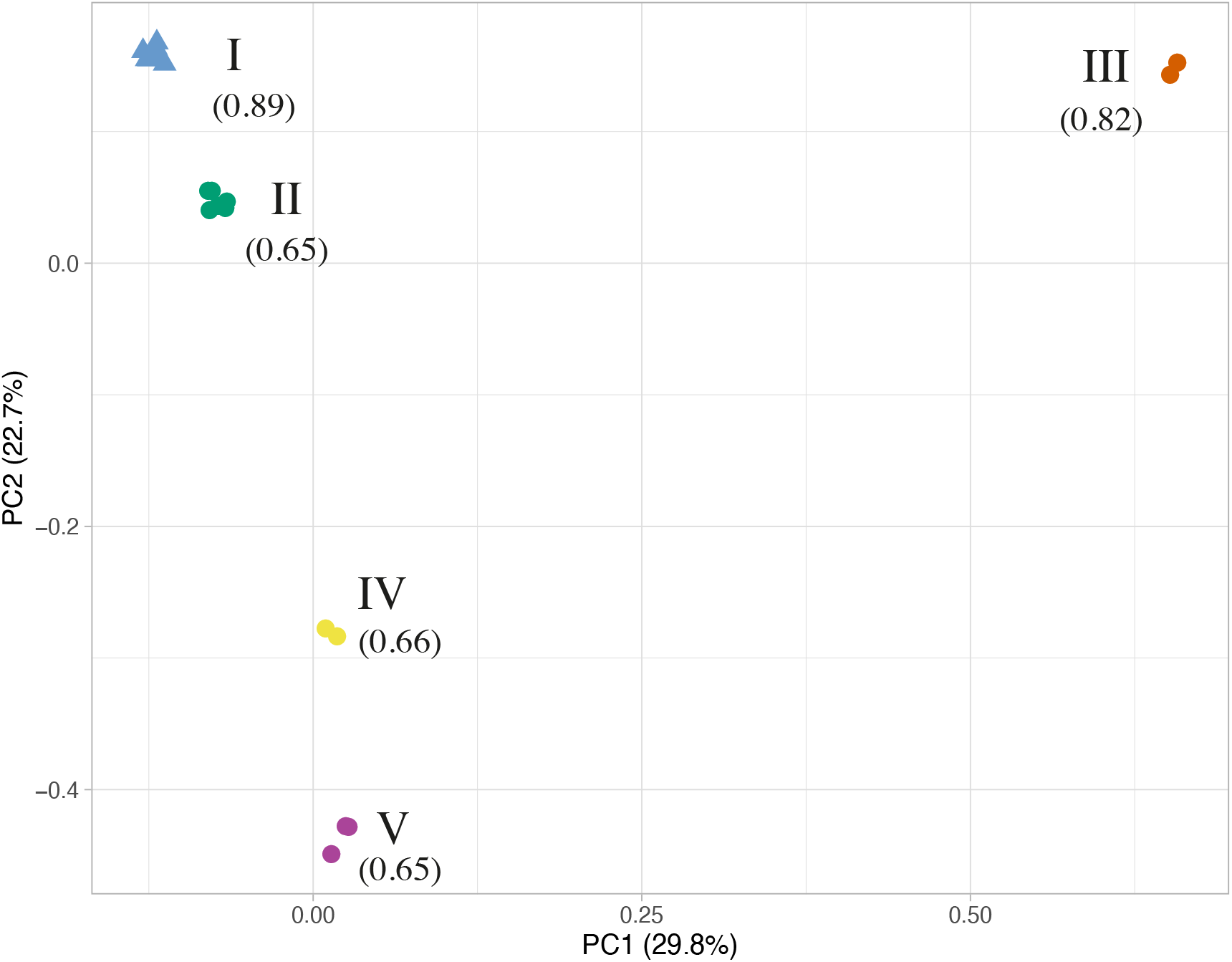
Principal component analysis based on nucleotide diversity among the 21 Y chromosomes (8190 polymorphic sites). The sensitive Y chromosomes are represented by a triangle while the resistant Ys are represented by a circle. Each Y chromosome is colored according to the cluster to which it belongs. Cluster numbers are indicated in roman numerals, with, in parentheses, the average female frequency produced over the Y chromosomes of the corresponding cluster in resistance tests.

None of the pairs of sensitive-resistant Y chromosomes originating from the same geographical area cluster together. This confirms that the resistant chromosomes did not emerge locally but rather arose from migration along with the driver. While the sensitive Y chromosomes (Egy07, Mor12 and Fra93) are older than their resistant counterparts, *i.e.* were present respectively in North Africa and Europe before the arrival of the Paris *SR* system, they appear to be very similar. Indeed, they group together into Cluster I. By contrast, the resistant chromosomes sampled more recently in those populations are more variable. The resistant Y chromosomes from Morocco (Mor16) and Egypt (Egy12) group together in Cluster II, while the resistant Y chromosome from France (Fra09) belongs to Cluster V. Thus, the resistant Y chromosomes from Morocco (Mor16) and Egypt (Egy12) appear to be closely related, whereas the resistant Y chromosome from France (Fra09) likely originated from a different migration path.

Finally, the two resistant Y chromosomes collected in Seychelles in 1981 (Sey81) and 2003 (Sey03) do not cluster together. They have very different resistance ability (Figure 1B) and each cluster with Y chromosomes showing a similar level of resistance (Figure 2). Because these Y chromosomes were collected more than 20 years apart, we cannot conclude if there was a polymorphism within the Seychelles islands with Y chromosomes from clusters II and III segregating in the population or if a migration event from continental Africa led to the secondary introduction of a more resistant Y chromosome in Seychelles islands.

### 3. Very little within-population variation of the Y chromosome

To get an insight into the intra-population polymorphism of the Y chromosome, we genotyped by PCR-sequencing 98 additional chromosomes from 22 populations. This first allowed us to validate 11 SNPs detected by our approaches and to look at within-population variation. We reconstructed a haplotype network (Supplementary Figure 2). It recapitulates the clusters identified by the PCA (Figure 2), except for cluster IV and V that grouped together as we do not have SNP distinguishing each cluster. Most of the time all the individuals collected in the same location at the same time carry the same haplotype (Supplementary Table2). Only one population from France (collected in 2014 and 2015) seems to be polymorphic, containing Y chromosomes from clusters I, II and V. We postulate that recurrent annual migrations occur in western Europe, in accordance with the observation that *D. simulans* cannot overwinter in temperate regions (Boulétreau-Merle, Fouillet, and Varaldi 2003; Machado et al. 2016). In addition, migration in *D. simulans* is likely largely human-aided and must have broadly amplified all over the world in the recent past, as suggested by the rapid invasion of the P-transposable element within the last decade (Hill, Schlötterer, et Betancourt 2016).

### 4. The resistant lineage predates the emergence of the driver

Previous work suggested that resistance to X^SR^ is ancestral to all *D. simulans* Y chromosomes because the species sister to *D. simulans* (*D. mauritiana* and *D. sechellia*) carry the CAA haplotype, which is associated with resistance in *D. simulans* (Helleu et al. 2019). We reexamined this hypothesis in the light of our new sequencing data. We sought to identify the corresponding allele carried by *D. melanogaster*, *D. mauritiana* and *D. sechellia* for the 8190 SNPs identified as polymorphic within *D. simulans*. We were able to recover 6138 SNPs in *D. mauritiana*, 6126 SNPs in *D. sechellia* and 3909 SNPs in *D. melanogaster*. Based on the 8190 SNPs we reconstructed the phylogeny of the Y chromosomes using a Maximum likelihood approach and *D. melanogaster* as the outgroup (Figure 3). The phylogeny defines 5 clades corresponding to the 5 clusters identified by the PCA (Figure 2). According to this tree it appears clear that the ancestral Y chromosome belongs to the resistant lineage. Indeed, the first divergent clade is cluster III, composed of two resistant Y chromosomes Sey81 and New91. Whereas the clade that includes all sensitive Y chromosomes (cluster I) appears to have a recent common ancestor nested within clusters of resistant Y chromosomes. The sensitive haplotype seems to diverge from the same ancestor as the resistant Y chromosome of cluster II.

**Figure 3.**
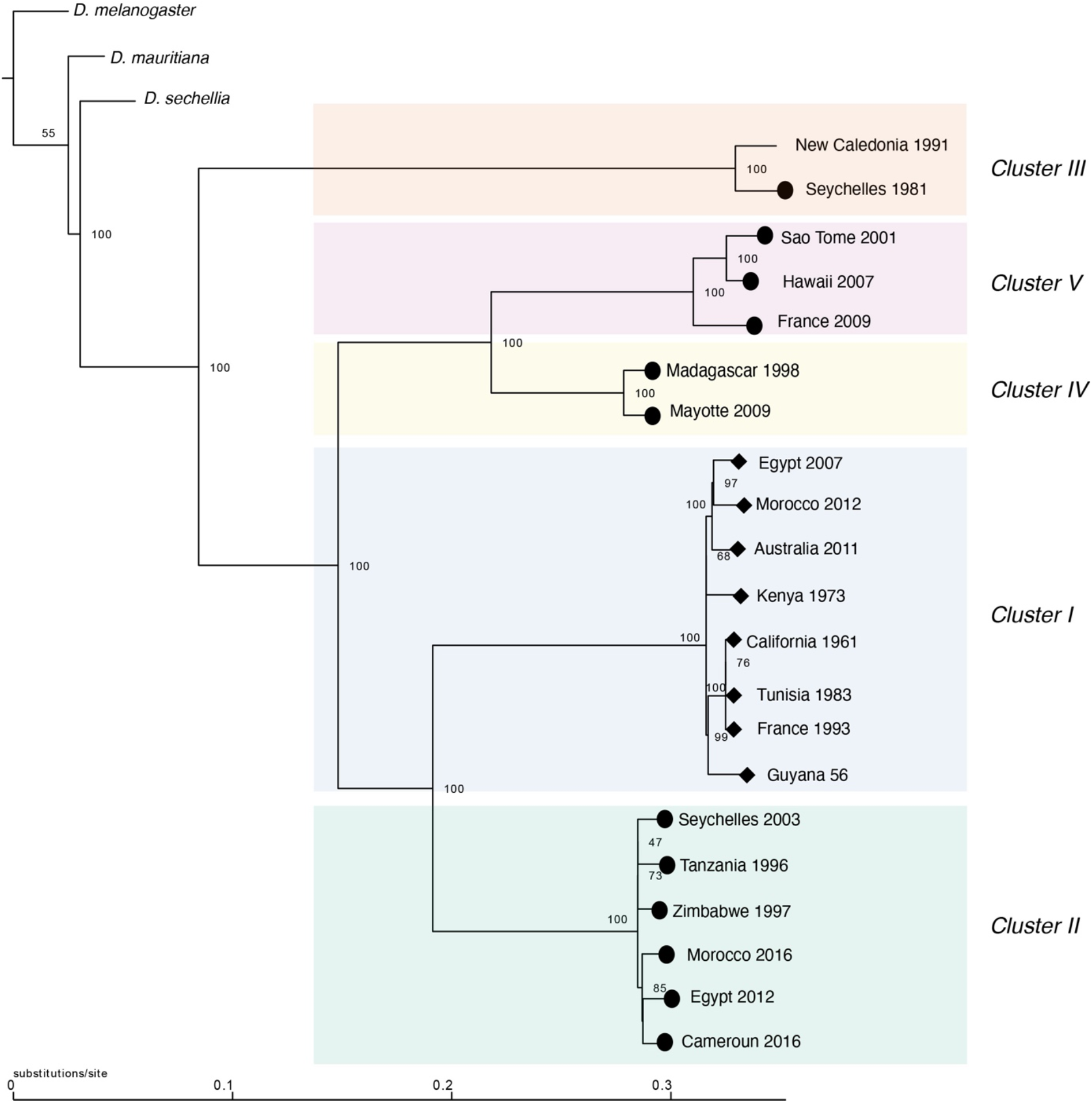
The inferred maximum likelihood phylogeny for the Y chromosome. *D. melanogaster* was used as the outgroup. Node labels indicate ultrafast bootstrap. The resistant Y chromosomes are indicated by a circle at the tip and sensitive Y chromosomes are indicated by diamond at the tip. Clusters are colored based on the cluster defined by the PCA analysis (Figure 2).

In addition, we focused on the 590 SNPs that correspond to fixed differences between the four clusters of resistant Y’s and the cluster of sensitive Y’s (Cluster I), to identify the allele carried by the sister species. While we were not able to recover all the SNPs, 97.4% of the recovered ones carry the resistant allele in *D. mauritiana*, 94.8% in *D. sechellia* and 93.7% in *D. melanogaster*.

Altogether, we conclude that the common ancestor of the *melanogaster* clade was carrying a Y chromosome closely related to the resistant lineage. Thus, while the sensitive lineage is the most common Y chromosome lineage found worldwide, it is a divergent lineage that seems to have a recent origin. This also means that the resistant Y chromosome lineage predates the emergence of the Paris *Sex Ratio* driver in *D. simulans*.

### 5. Gene copy numbers appear stable among Y chromosomes

To assess whether some components of the Y chromosomes are associated with the resistance phenotype, we first estimated the exon copy number for each of the 11 canonical Y-linked genes (Figure 4A). Our estimates are in agreement with the previous estimates from the reference genome (Chang et al. 2022). Most of the canonical genes are in single copy and do not exhibit variation between Y chromosomes (Figure 4A). ARY was previously described as a multi copy gene with 4 copies on the reference genome; here we estimated between 4 and 6 copies of ARY among our 21 Y chromosomes. Similarly, *kl2* exons 8 to 12 are known to be duplicated, with more than 15 copies for exon 10 for example. They are also the most variable among the 21 Y chromosomes, with *kl2*-exon10 showing between 10 and 17 copies. *kl3*-exon16 shows an interesting variation: while most of the Y chromosomes have 3 copies, the 3 Y chromosomes that belong to Cluster V (Haw07, Sao01, Fra09) show a high duplication rate (between 7 and 11 copies) (Figure 4A).

**Figure 4:**
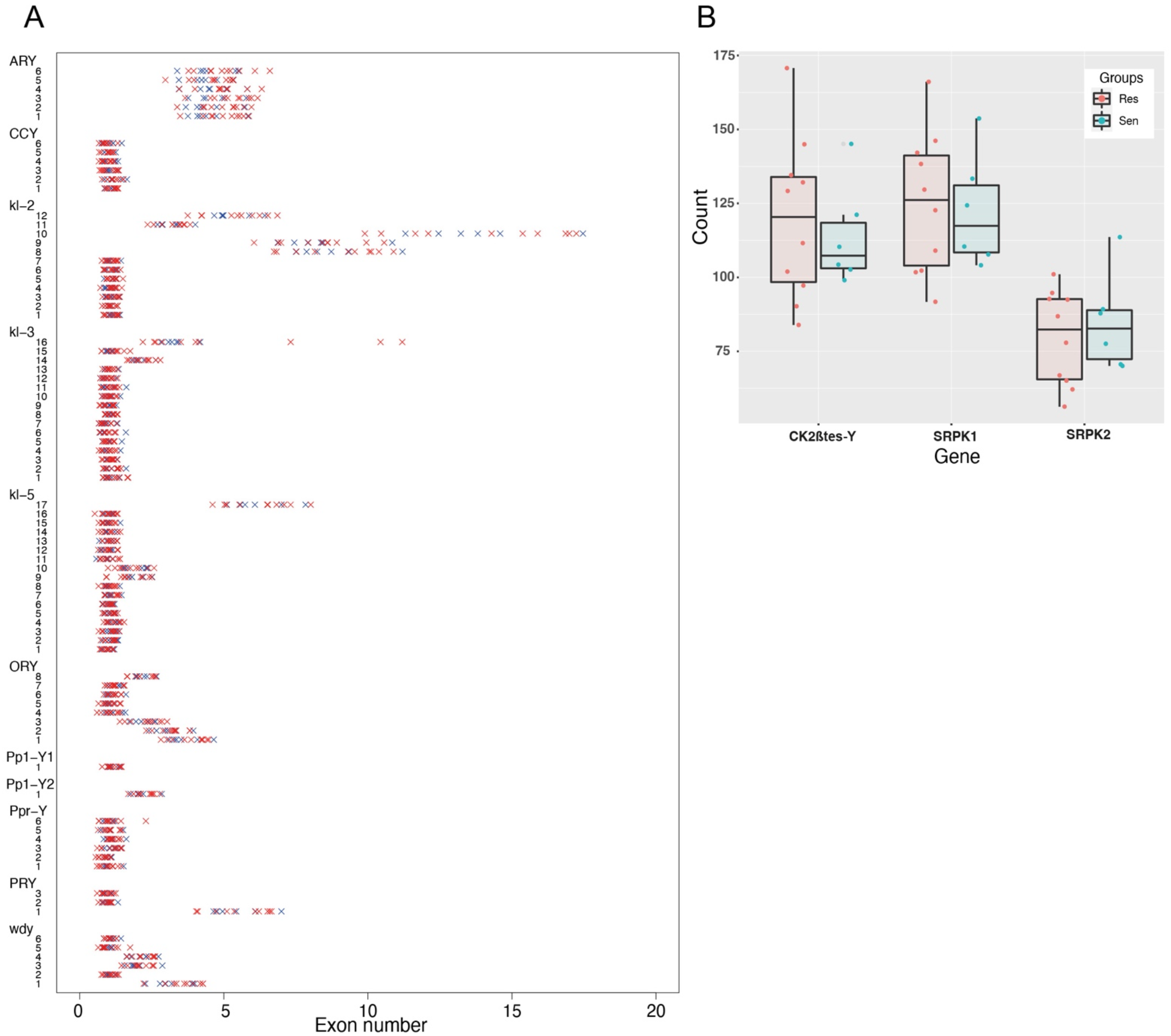
A. Strip chart representing the exon numbers for each Y-associated gene. Each line corresponds to a specific exon of a specific gene. Red crosses correspond to resistant Y chromosomes and blue crosses to the sensitive Y chromosomes. B. Boxplot representing the exon numbers for the two Y-linked multicopy gene families, *CK2βtes-Y* and *Lhk*, among resistant (Res) and sensitive (Sen) Y’s.

In addition to the 11 canonical genes, two ampliconic genes have been described on the Y chromosome of *D. simulans* : *CK2βtes-Y* and *Lhk* (Chang et al. 2022). Accordingly, both exhibit variation in exon copy number among our 21 Y chromosomes. We estimate between 83 and 170 copies of *CK2βtes-Y*, between 91 and 156 copies of *Lhk-*exon1 and between 56 and 113 copies of *Lhk*-exon2, depending on the Y (Figure 4B).

In general, only exons that are multi-copy appear to be variable in terms of copy number. Importantly, we did not find any evidence for an association between the exon copy number and the resistance phenotype (Figure 4 A and B). It thus appears unlikely that the variation in resistance is due to a variation in exon copy number.

### 6. Very low Y-linked variation in transposable element abundance between populations

Using the raw Illumina reads we then estimated the abundance of transposable elements and complex satellites in 16 iso-Y lines (6 sensitive and 10 resistant; see materials and methods). Because our Y chromosomes have been isolated in the same genetic background, we assumed that variation in the abundance of repeats between iso-Y lines is associated with the Y chromosome. To investigate whether variation in Y-linked complex repeat abundance shows the same clustering as Y-linked nucleotide polymorphism, we performed a PCA analysis (Figure 5A). Globally the complex repeat abundance does not replicate the clusters defined by SNPs in Figure 2. The sensitive Y chromosomes (Cluster I) show a diffuse distribution that partially overlaps with some resistant Y chromosomes. Similarly, the distributions of resistant Y chromosomes from clusters III, IV and V largely overlap. Only the Y resistant chromosomes from cluster II fully separates from all the others.

**Figure 5:**
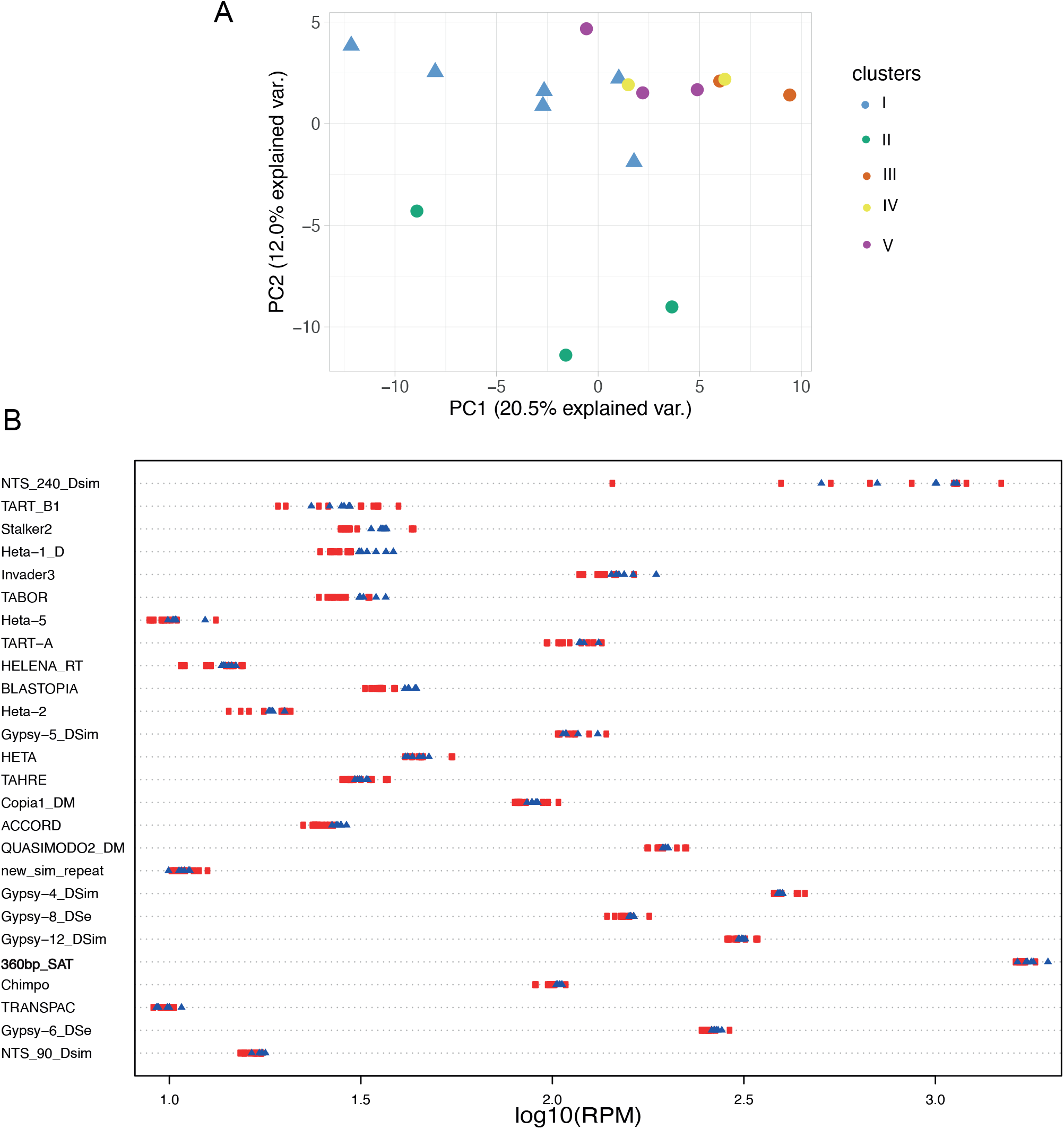
A. Principal Component Analyses (PCA) based on the complex satellite abundance. Sensitive Y chromosomes are represented by a triangle while the resistant Ys are represented by a circle. Each Y chromosome is colored based on the clusters defined previously using the nucleotide diversity (Figure 2). B. Strip chart representing the top 25 most variable complex satellites (based on coefficient of variation), among which the *360 bp* satellite (in bold) is indicative of the background noise. To be able to distinguish the differences between strains, Reads Per Million (RPM) counts have been transformed in log10. Each dot corresponds to a Y chromosome. The Y resistant chromosomes are represented by a red circle while the sensitive Y chromosomes are shown with a blue triangle.

Next, we tested whether the phenotypic variations observed among the Y chromosomes were associated with variations in gross repeat content. We thus performed a Wilcoxon test to assess whether some transposable elements or complex satellites might be differentially enriched between sensitive and resistant Y chromosomes. None of these repeats show significant variation (Supplementary Table3), indicating that the abundance in complex repeats is not associated with the resistance phenotype. More generally, it appears that most complex repeats exhibit very low to no variation in abundance between iso-Y lines (Supplementary Table3). We calculated the coefficient of variation for each complex repeat and plotted the abundance for the Top25 most variable ones (Figure 5B). The NTS-240 element is the only complex repeat that shows substantial variation, with more than 10-fold variation in abundance between Y lines (Figure 5B). While lower in abundance, the telomeric transposable elements (Het-A, TART and TAHRE) are also among the most variable repeats (Figure 5B).This is in agreement with the existence of those elements in various regions on the Y chromosome (Agudo et al. 1999).

However overall, the majority of these Top 25 most variable repeats show relatively low variation between iso-Y lines. It is important to note that the abundance represented here is the whole genome abundance. Thus, complex repeats that are not associated with the Y chromosome are expected to show low to no variation, the level of which could reflect that of background noise. This is the case for the *360bp* satellite, which is a complex satellite in the 1.688 family, highly abundant on chromosome II and III in *D. simulans* (Sproul et al. 2020; de Lima, Hanlon, et Gerton 2020) and not associated with the reference Y chromosome (Chang et al. 2022). The *360-bp* satellite shows 1.8-fold variation among iso-Y lines, and we therefore assume that repeat variations at or below this threshold likely represent background noise (Figure 5B).

### 7. Simple satellite enrichment reveals candidates for the resistance

Finally, we compared the simple satellite composition between the same 16 iso-Y lines (6 sensitives and 10 resistants). Simple repeats represent over 5% of *D. simulans* genome and are a major component of the Y chromosome (Wei et al. 2018). Using the k-seek pipeline (Wei et al. 2014), we identified 177 simple satellites, including the most abundant kmers previously described to be highly enriched in *D. simulans* (Wei et al. 2018) (Supplementary Table4). Interestingly, we found that simple satellite abundance recapitulates the structure between iso-Y lines defined by the SNP calling (Figure 6A vs Figure 2).

**Figure 6.**
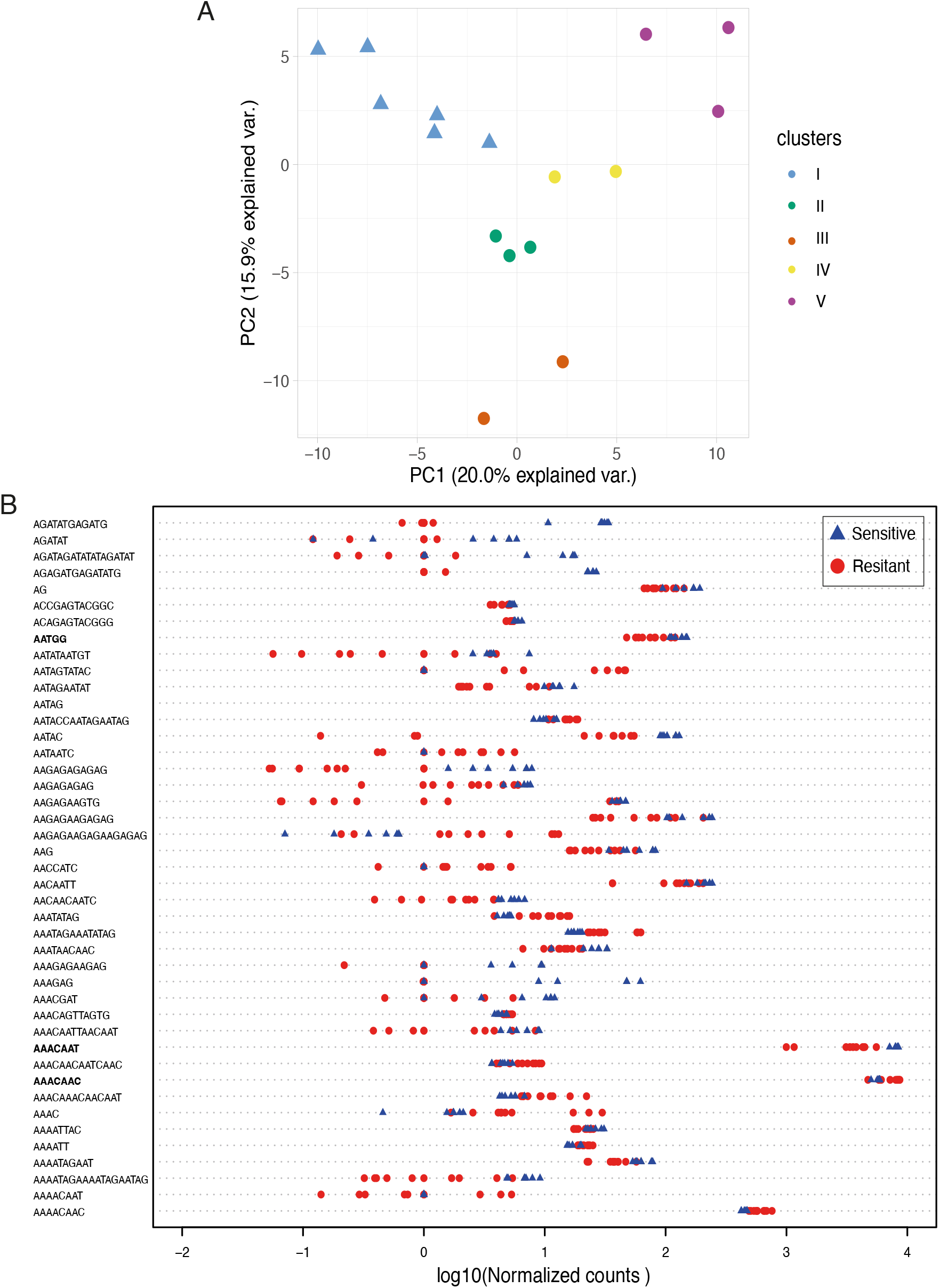
A. Principal component analyses (PCA) based on the simple satellite abundance. Each Y chromosome is colored according to the cluster to which it belongs (defined by nucleotide diversity, Figure 2B). Sensitive Y chromosomes are represented by triangles while the resistant ones are represented by circles. C. Strip chart of the 43 simple satellites differentially enriched (p.value<0.05) between the resistant Y’s (red circles) and the sensitive Y’s (blue triangles). Normalized counts were transformed in log10.

Among the 177 satellites, 43 appear differently enriched on resistant versus sensitive Y chromosomes (Wilcoxon test, p.value<0.05) (Figure 6B). About two third of them (28/43) are more enriched among sensitive Y chromosomes (Figure 6B, Supplementary Figure 3). Because we observed variation in resistance among the former group (Figure 1B), we performed a Spearman correlation test between the abundance of each of the 177 satellites and the sex ratio observed in the resistance tests for the 10 resistant Ys plus the reference sensitive Y (Tunisia 1983), (Supplementary Figure 4). Based on their abundance and correlation with the phenotype and on their location (enriched or specific to the Y chromosome), we identified a set of potential candidate satellite that may be involved in the resistance phenotype.

Among the satellites more abundant on sensitive Y chromosomes, four of them, known to be associated to the Y chromosome (Wei et al. 2018), are particularly interesting. The first one, (AAACAAT)n is among the most abundant satellites and is 2-fold more enriched in the sensitive lineages than in the resistant lineages (Figure 7A, Wilcoxon test pvalue<2e-04). The other three, (AATGG)n, (AAGAGAAGAGAG)n and (AACAATT)n, are less abundant, but the sensitive Y chromosomes are also significantly more enriched than the resistant ones (Wilcoxon test, pvalue<0.05). None of these satellites show significant correlation with the strength of the resistance phenotype (Supplementary Figure 4). We performed FISH on mitotic chromosomes for a subset of our iso-Y lines from each cluster, with probes for (AAACAAT)n, (AATGG)n (Figure 7B, Supplementary Figure 5) and (AAGAGAAGAGAG)n and (AACAATT)n (Supplementary Figure 6). We confirmed that (AAACAAT)n, (AATGG)n and (AACAATT)n are specific to the Y chromosomes, at least at the level FISH can detect. With regard to (AAGAGAAGAGAG)n, a signal was observed on the Y chromosomes and also on the X chromosome (originating from the ST8 line, see Mat and Met) (Supplementary Figure 6). We observed a strong signal with AAACAAT for all the Y chromosomes tested, which is consistent with its high copy number. The signal for AATGG was clearly lower as expected from its lower abundance (Figure 7A). While we detected a AATGG signal for all the sensitive Y chromosomes, the signal was low or absent for the resistant Y chromosomes (Figure 7B, Supplementary Figure 5). Once again, this is consistent with our estimates that there are fewer copies on the resistant Y chromosomes. In addition to quantitative variation, we observed structural variations associated with the four satellites (Figure 7B, Supplementary Figure 5-6). In most of the cases they seem located at or close to the tip of the Y chromosomes long arm, while they appear closer to the centromere on the resistant chromosomes from cluster II (Figure7B, Supplementary Figure 5-6).

**Figure 7.**
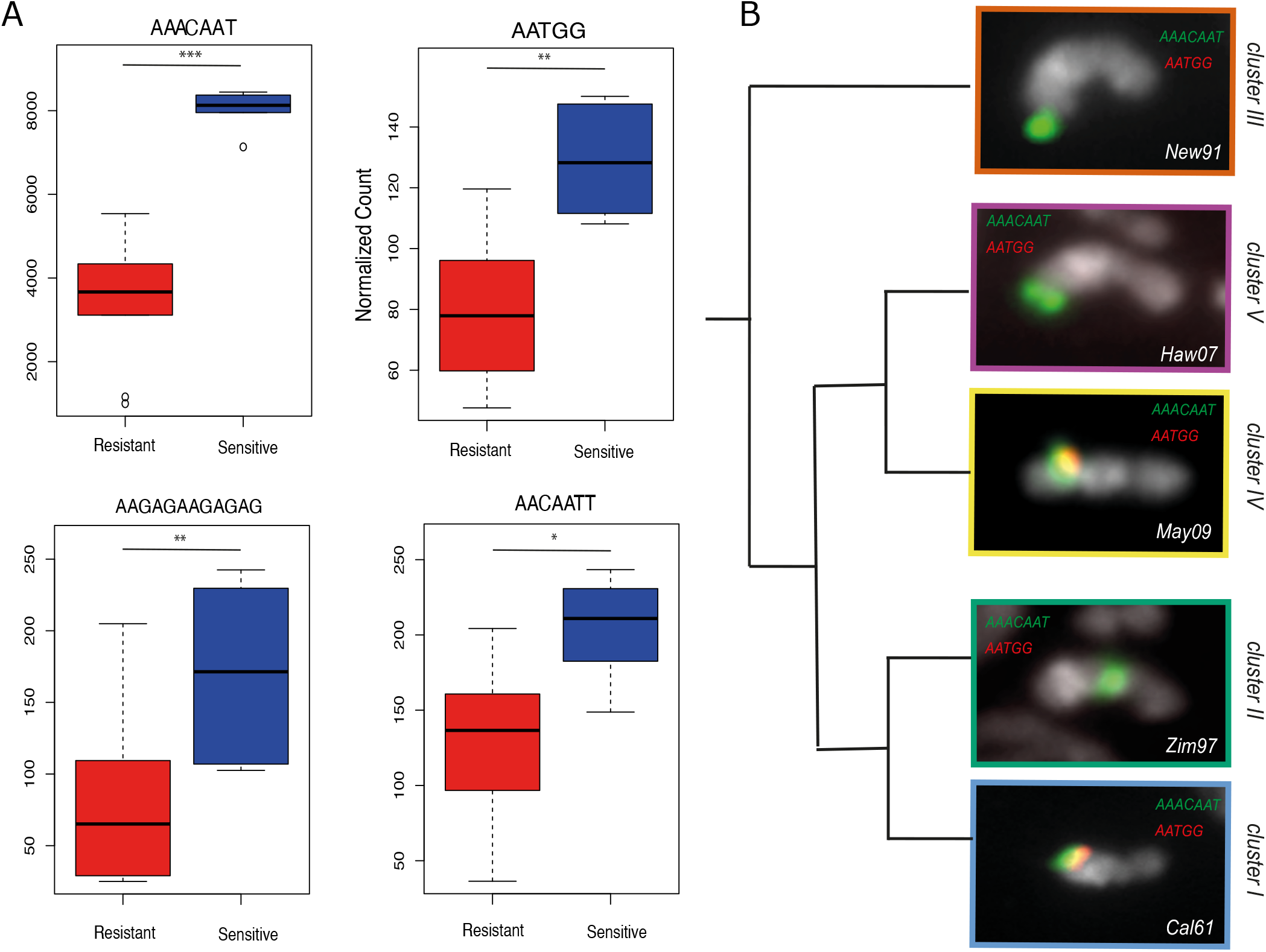
A. The box plots represent the abundance of (AATGG)n, (AAACAAT)n, (AAGAGAAGAGAG)n and (AATGG)n among resistant Ys (red) and sensitive Ys (blue). B. FISH on mitotic chromosomes from larval brains using (AACAAT)_4_ (green) and (AATGG)_6_ (red) probes. A single Y chromosome from each cluster is displayed. The whole karyotype FISH on the other iso-Y lines are shown in Supplementary Figure 5.

Finally, two additional satellites showing an opposite pattern came to our attention, (AAACAAC)n and (AACAATC)n. While they show no or low differential enrichment between the sensitive and resistant Y chromosomes (Figure 8A-B), they show a strong positive correlation between abundance and the resistance level (Figure 8C-D, Spearman test, p.value<0.01). The most resistant Y chromosomes are also the most enriched for (AAACAAC)n and (AACAATC)n. Both satellites are known to be associated with the Y chromosome (Wei et al 2018), (AACAATC)n is even Y-specific. Using FISH on mitotic chromosomes, we show that the signal of both satellites is specific to the Y chromosome (Figure 8E). And seems to have a localization similar to the previous candidates (Figure 7B and 8E). (AAACAAC)n and (AACAATC)n are located close to the tip of the long arm of the Y chromosome except for the Y chromosomes from cluster II, where they appear closer to the centromere (Figure 8E, Supplementary Figure 7).

**Figure 8:**
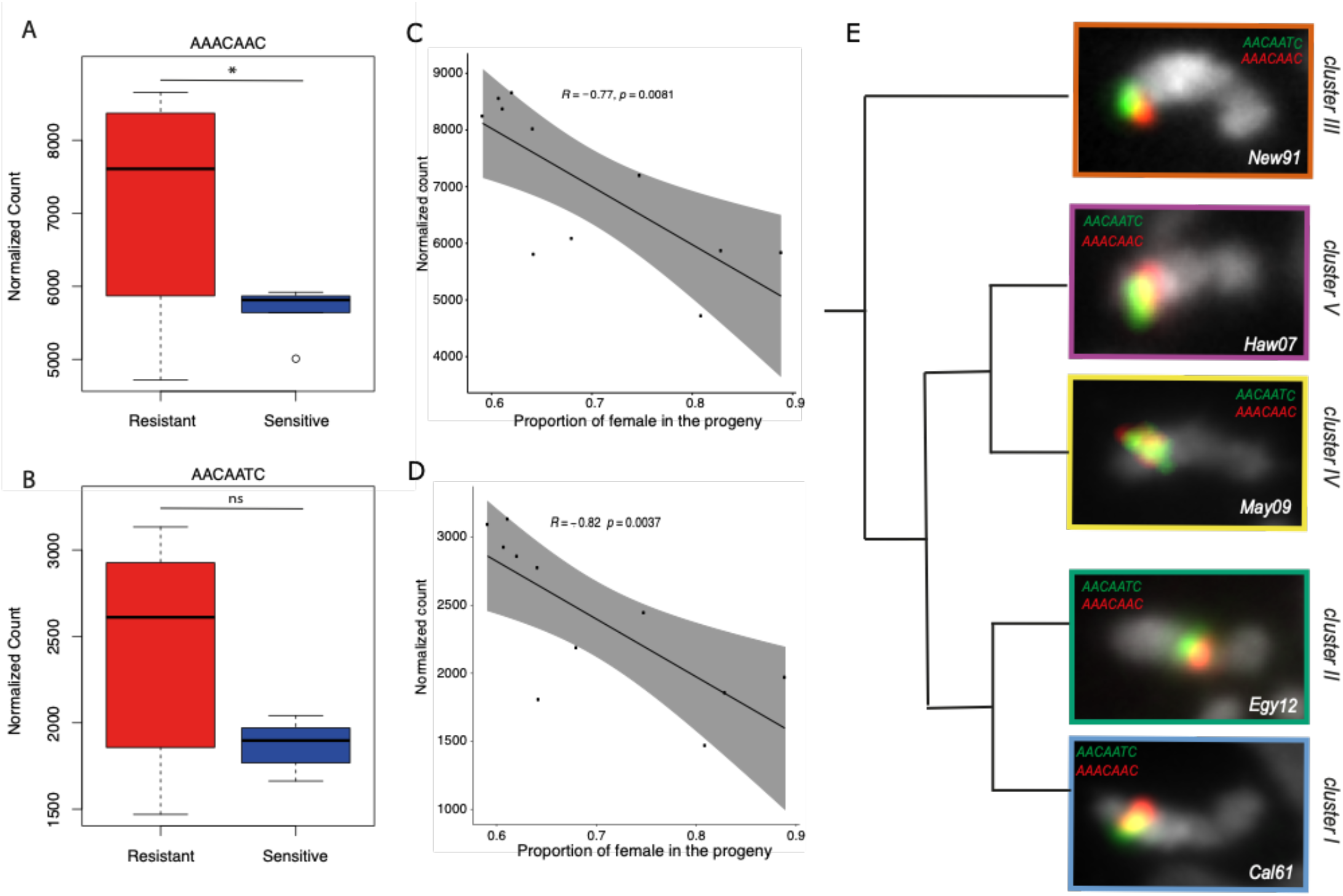
Abundance of (AAACAAC)n (A) and (AACAATC)n (B) satellites. The box plots represent their abundance in each group of Y chromosomes (resistant in red and sensitive in blue). C and D. The dot plots show the correlation between the satellite abundance (y axis) and the proportion of females in the progeny (x-axis), inversely proportional to the resistance. Correlation was calculated based on a Spearman correlation test. E. FISH on mitotic chromosomes from larval brains using (AACAATC)_4_ (green) and (AAACAAC)_4_ (red) probes. A single Y chromosome from each cluster is displayed. The whole karyotype FISH on the other iso-Y lines are shown in Supplementary Figure 7.

To further validate the accuracy of our abundance estimates by k-seek, we quantified the abundance of AAACAAT, AAACAAC and AACAATC using slot blot. We found a positive correlation (Pearson correlation test, pvalue <0.05) between the abundance inferred by k-seek and the blot quantifications (Supplementary Figure 8).

## DISCUSSION

Y-linked nucleotide diversity in both *Drosophila melanogaster* and *Drosophila simulans* is known to be very low compared to the rest of the genome (Larracuente et Clark 2013; Helleu et al. 2019). Those previous estimates were done using genic sequences, which limits the estimation to a very small portion of the Y chromosome. Here, taking into account the assembled part of the chromosome (around 13 Mb) we found that the Y-linked nucleotide diversity in *D. simulans* is about 15-fold lower than the estimate in *D*. *melanogaster* (π was estimated ranging from 0.00266 to 0.01696 in Larracuente et Clark 2013, here π = 0.0001808). While the level of nucleotide variation among the Y chromosomes examined here is very low, it reflects the evolutionary history of the species.

### Evolutionary history of the Y chromosome related to the demographic history of D. simulans

#### The ancestral origin of the Resistance

*D. simulans* is thought to originate from Madagascar or the surrounding area (Lachaise et Silvain, 2004.; Dean et Ballard 2004). We hypothesize that this ancestral population was carrying a drive resistant Y chromosome (Figure 6A). Indeed, according to the phylogeny (Figure 3), the sensitive Y chromosomes are derived recently in *D. simulans* from resistant Y chromosomes, suggesting that the ancestral Y chromosome in *D. simulans* was of the resistant type. The ancestry of the resistant lineage does not necessarily mean that the resistant phenotype is ancestral. However, based on our phylogeny it is most parsimonious that the sensitive lineage (Cluster I) corresponds to a secondary loss of resistance to Paris SR. Consistent with this scenario, previous work suggested that the Y chromosome of *D. sechellia* is resistant to the Paris drivers (Tao et al. 2007; Rice 2014).

#### Spread of *D.simulan*s in Seychelles and Continental Africa

During a first expansion event, *D. simulans* is thought to have independently colonized the Seychelles islands and the African continent (Lachaise et Silvain, 2004) (Figure 6A). Our data support this scenario. Indeed, most of the Y chromosomes from continental Africa belong to Cluster II while the two chromosomes from Seychelles and New Caledonia belong to Cluster III (Figure 2). The phylogeny agrees with the early divergence of these two chromosomes. Not only do they appear genetically different but they also have a different phenotype. Indeed, the Y chromosomes from Seychelles (Sey81) and New Caledonia (New91) appear less resistant than the other resistant Y chromosomes (Figure 1B).

This scenario for the demographic history of *D. simulans* was supported by the distribution of the mitochondrial (mt) haplotype. In *Drosophila simulans* there are three different mt haplotypes : *si*I, *si*II and *si*III (Solignac et Monnerot 1986). The *si*II haplotype is found worldwide. In Madagascar and La Reunion, populations appear polymorphic for *si*III and *si*II. The last haplotype, *si*I, is specific to the Indo-Pacific islands (Seychelles, New Caledonia and Hawaii). We controlled for the mitochondrial haplotype of our strain. As expected most of them carry the *si*II haplotype with the exception of the strains from Hawaii, New Caledonia and Seychelles that carry the *si*I haplotype, and Mayotte that carries the *si*III haplotype (Supplementary Table5).

#### Emergence and spread of the sensitive lineage

The phylogeny indicates that the sensitive Y chromosomes (Cluster I) and resistant Y chromosomes from Continental Africa (Cluster II) have a recent common ancestor (Figure 3). This suggests that the sensitive lineage emerged in Continental Africa and started to replace resistant Y chromosomes (Figure 9B). This hypothesis is supported by the Ken73 Y chromosome that comes from flies collected in Kenya in 1973. The Ken73 Y chromosome belongs to Cluster I and appears similar to the sensitive Y chromosomes found across the world. The Y chromosome that has then spread out-of-Africa is the sensitive Y chromosome, which became the major Y chromosome across the species range (Helleu et al. 2019) (Figure 1A, Figure 6C). The low nucleotide diversity (>97% similarity, Table 1) within the sensitive Y lineage suggests that this event probably arose very recently. Indeed it has been shown that the expansion of *D. simulans* is more recent than that of *D. melanogaster*, likely 6500-5000 years in Eurasia (Lachaise et Silvain, 2004) and 500 years in North America (Wall, Andolfatto, et Przeworski 2002).

**Figure 9.**
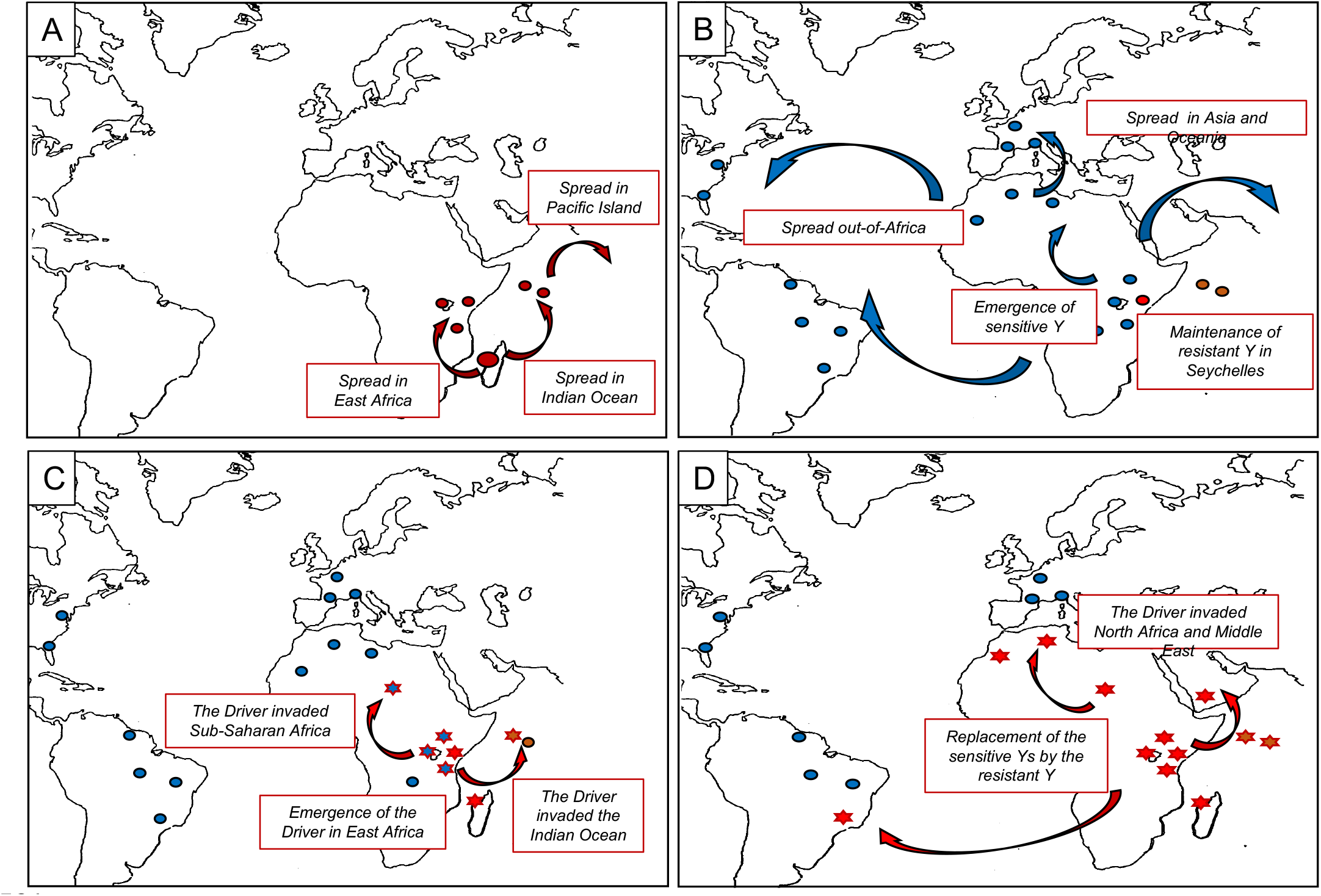
Proposed scenario for the demographic history of *D. simulans* in relation to the Paris *SR* system. A. Expansion of *D. simulans* out of its area of origin, toward the Seychelles and the African continent (Lachaise et Silvain, 2004.; Dean et Ballard 2004). The ancestral Y chromosome belongs to the resistant lineage, here represented by a red circle. B. Emergence of the sensitive lineage in subsaharan Africa and diffusion out-of-Africa. Here sensitive Y chromosomes are represented by a blue circle. C. Emergence of the Paris *SR* system in East-Africa (Bastide et al. 2011). Populations where the system is detected are represented by a star. D. Spread of the Paris *SR* system and replacement of the sensitive Y chromosome by resistant Y chromosome (Helleu et al. 2019).

**Table 1:** Summary of the number of shared SNPs per cluster.

| Groups | Sensitive<br>(Cluster I) | Resistant<br>(all) | Resistant<br>(Cluster II) | Resistant<br>(Cluster III) | Resistant<br>(Cluster IV) | Resistant<br>(Cluster V) |
| --- | --- | --- | --- | --- | --- | --- |
| N° shared<br>SNP | 6985 | 1123 | 6191 | 7488 | 7644 | 6704 |
| Average %<br>of identity | 97.77 | 76.89 | 97.78 | 96.13 | 97.58 | 95.77 |

Why has the sensitive Y chromosome previously superseded the ancestral resistant Y chromosome? Multiple non-exclusive factors could have favored the spread of Y sensitive lineage. First, the emergence of an advantageous mutation may have led to a strong positive selection on sensitive Y chromosomes. A selective advantage associated with a Y chromosome can quickly lead to its fixation into the population (Clark 1987). The Y sensitive chromosomes might improve the fitness of their carrier over the resistant Y chromosome, they can increase the fertility or favor the adaptation to a new environment. Second, the demographic history of the species could also explain the low level of polymorphism of the sensitive Y chromosomes (Irvin et al. 1998; Hamblin et Veuille 1999). Indeed, effects such as migration and bottlenecks could have facilitated the spread of the sensitive lineage and reduced its effective population size. Finally, the last hypothesis involves recurrent genetic conflict. In *D. simulans,* there are at least 3 independent *SR* systems: Paris, Winters and Durham. The Winters and Durham SR systems are cryptic and presumably much older than the Paris system (Meiklejohn et al. 2018). Thus, *SR* systems appear common in this species (Muirhead et Presgraves 2021; Vedanayagam, Lin, et Lai 2021). The intense resulting genetic conflicts between X and Y chromosomes have an important impact on sex chromosome evolution. We hypothesize that the presence of an X-linked driver in *D. simulans*, before the emergence of the Paris *SR* system, could have driven the spread of some Y chromosomes belonging to the lineage sensitive to the Paris driver. Some of the Y chromosomes sensitive to the Paris driver could have been resistant to this more ancient driver. A quick replacement of sensitive by resistant Y chromosomes might have occurred in the past but triggered by another driver. This more ancient driver would have favored the spread of the sensitive Y chromosome lineage before becoming cryptic or disappearing. Previous work shows large variation in the resistance ability of the Y chromosomes to the Winters driver (Branco et al. 2013).Testing the same set of Y chromosomes in the three *sex-ratio* systems might help understanding the cause of the spread of the sensitive Y chromosomes.

#### Emergence of the Paris system and replacement of the sensitive by the resistant Y lineage

The Paris system likely emerged in East Africa less than 500 years ago and quickly spread in the Indian Ocean Island and Sub-Saharan Africa (Bastide et al. 2011). During the last two decades we observed the invasion of the system in North Africa, Europe and the Middle East (Bastide et al. 2013; Helleu et al. 2019). Due to their transmission advantage the resistant Y chromosomes were strongly selected and quickly spread along with the driver, replacing the Y sensitive chromosomes (Helleu et al. 2019) (Figure 9D).

Our approach allows us to follow the migration path of the Paris SR system. For example, the resistant Y chromosomes from Egypt, Morocco and France segregate in different clusters, indicating that the Paris SR system has migrated independently to North Africa and Europe. Similarly, the two resistant Y chromosomes from Seychelles - which have been collected more than 20 years apart - belong to two different clusters. The Y chromosome collected in 2003 (Sey03, cluster II) likely results from migration from continental Africa to the Seychelles island around that time. In addition, it appears to be more resistant than the one collected in 1981 (Sey81, cluster V), which could explain this replacement.

#### Genetic basis of the resistance

Beyond the evolutionary history of the Y chromosome, we also provide important insight into the genetic basis of the resistance to Paris X^SR^. Based on its large phenotypic variation we hypothesize that the resistance ability might be associated with variation in the repeat content on the Y chromosome. We did not find any association between the resistance and the gene or transposable element abundance on the Y chromosomes, they even appear quite stable. In contrast, we do find a strong association with the simple satellite composition (Figure 6A) and identified several candidates that could be involved in resistance: (AAACAAT)n, (AATGG)n, (AAGAGAAGAGAG)n and (AACAATT)n, (Figure 7A) or (AAACAAC)n and (AACAATC)n (Figure 8A-B). (AAACAAT)n, (AATGG)n, (AAGAGAAGAGAG)n and (AACAATT)n are known to be enriched or specific to the Y chromosome in *D. simulans* (Wei et al. 2018) (Supplementary Figure 5-6) and exhibit a lower copy number on the resistant Y chromosome (Figure 7A). In addition to the copy number variation, (AAACAAT)n, (AATGG)n appear to be associated with structural variation among resistant Y chromosomes (Figure 7B, Supplementary Figure 5) (Helleu et al. 2019), which may also contribute to variation in the phenotype. The two other candidates, (AAACAAC)n and (AACAATC)n, are also enriched on the Y chromosome (Wei et al 2018) but, on the opposite, their abundance positively correlates with their resistance ability (Figure 8). This would suggest that those satellites could confer the ability to resist to the driver. One of the drivers of the Paris *SR* system has been identified as *HP1D2*, a member of the heterochromatin protein gene family, known to be involved in heterochromatin regulation. We have previously shown that HP1D2 is specifically expressed in spermatogonia and is enriched on the Y chromosome (Helleu et al. 2016). In addition, the drive is due to a loss of function of *HP1D2*. We thus hypothesize that the drive is associated with a mis-regulation of the heterochromatin region of the Y chromosome. While we do not know the precise target of HP1D2, those simple satellites are possible candidates. Correlations between the target copy number and the strength of the drive has been shown in other meiotic drive systems. For example, in the *SD* system of *D. melanogaster*, the *Rsp* satellite is the target of the driver (Hartl 1974). Variation in *Rsp* copy number is related to variation in the sensitivity to the driver (Lyttle 1989). Chromosomes with a higher copy number of *Rsp* are more sensitive to the driver. Similarly, particular repeats may be involved in the resistance to the Paris drivers and possibly be their target. Four of the simple satellite candidates we identified as associated with drive sensitivity of the Y chromosome, follow this model (Figure 7A). Three of them, (AATGG)n, (AAACAAT)n and (AAGAGGAAGAGAG)n, appear absent from the *D. sechellia* Y chromosome (Wei et al 2018). This is consistent with the finding that the *D. sechellia* Y chromosome is resistant to the Paris driver. Variation in resistance could be due to variation in their abundance and/or in their localization. Considering that the distortion is due to a misregulation of heterochromatic sequences, the localization of those sequences on the Y chromosome could be more or less critical for chromosome segregation. Also, we cannot exclude that the resistance phenotype is multifactorial, or that the genetic basis of resistance is different between Y chromosome lineages.

To conclude, our study sheds light on the evolutionary history of the Y chromosome in *D. simulans*. We show that resistant Y chromosomes likely predated the emergence of the Paris *SR* system and that sensitive Y chromosomes have spread through the world in the recent past. The Paris system is particularly young, at least in its present form, since the segmental duplication which is involved in driving likely emerged less than 500 years ago (Bastide et al. 2013). While its impact on the recent evolutionary history of the Y chromosome is clearly strong, it cannot explain on its own the whole pattern of variation we observed here. More likely, the Y chromosome was recurrently involved in genetic conflicts, which resulted in multiple selective sweeps. Furthermore, we also provide new insights on the genetic basis of Paris drive resistance. Knowing that simple satellites are likely involved should help us to move further in the understanding of the molecular mechanism underlying the Paris *SR* system.

## MATERIALS AND METHODS

### Fly stocks

Except for Ken73, all iso-Y lines (derived from a single male) carry the same standard genetic background (X^ST^ A^ST^) from the ST8 reference strain (Montchamp-Moreau et Cazemajor 2002). The crossing procedure used to isolate the Y chromosomes is presented in(Montchamp-Moreau, Ginhoux, et Atlan 2001). The fourth chromosome was not controlled. The place and date of collection of the strains from which the Y chromosomes studied originate are given in Supplementary Table5.

ST8 is the reference standard strain in the Paris *SR* system. It has been collected in Tunisia (1983) and is free of driver and suppressor elements (Montchamp-Moreau et Cazemajor 2002). The *ST8/C(1)RM,yw* line is derived from the reference standard line ST8. Females carry a compound X chromosome *C(1)RM,yw* and males carry the X^ST8^ from the ST8 line. The Y chromosome and the autosomes are from the ST8 line.

The *ST8;Cy;Delta/Ubx* line carry the sex chromosomes from the ST8 line and dominant markers on chromosomes II (^Cy^) and III (^Ubx/Delta^).

The X^SR3^ line is used to test the resistance ability of the Y chromosome. The males carry an X^SR^ from the reference SR strain (collected in Seychelles in 1981, Cazemajor, Landré, et Montchamp-Moreau 1997) and the females carry the compound Xchromosome*C(1)RM,y,w*. The Y chromosome and the autosomes come from the ST8 line.

For simplicity each iso-Y line name is coded as follows : the 3 first letters of their location + last two digits of the collection year (for instance Morocco 2012 is Mor12).

### Resistance test

Each Y chromosome from the CAA lineage, and the standard Y^ST8^ have been tested for their resistance ability against the same reference X^SR3^. The crossing procedure is detailed in Montchamp-Moreau, Ginhoux, et Atlan 2001. Briefly, males carrying the Y chromosome to be tested (Y^pop^) were mass-mated with *ST8/C(1)RM,yw* females. Three F1 females were individually mated with X^SR3^ males, thus producing F2 males carrying both the X^SR3^ and Y^pop^ and standard autosomal background. Five males from each F2 progeny were individually mated, each with 2 ST8 females (so a total of 15 F2 males tested for each Y^pop^). Only individual F3 progenies consisting of at least 50 flies were considered.

Before statistical analyses, the data were transformed in *arcsin(*√𝑞), where *q* corresponds to the proportion of females in the progeny. To testforwithin-strain variation in Y resistance, we performed a Kurskal Wallis test between groups of F2 progenies (from the same F1 cross). All the iso-Y lines appear homogeneous (Supplementary Table1). We performed a Wilcoxon test to determine whether those lineswere significantly different from ST8. They all appear significantly more resistant than ST8 except Sey81 and New91 (Supplementary Table1). Finally, we performed a Tukey HSD post-hoc test on the anova model to determine groups (Figure 1B). All statistical analyses were conducted in R base(R Core Team 2022). The R script used can be found on github: Figure1/resistance.R.

### DNA extraction and sequencing

DNA extraction and sequencing was performed in two batches. In batch1 we sequenced: Tun83, Egy07, Ken73, Zim97, Egy12, Tan96 and Sey81. In batch2 we sequenced: Tun83, Fra93, Aus11, Mor12, Cal61, Guy56, Sey81, May09, Fra09, Mor16, Sao01, Cam16, New91, Haw07, Mad98, Sey03. Tun83 was sequenced in both batches to control for potential technical bias. Sey81 was sequenced in the first batch in its original genomic background (from Seychelles 1981) and then the Y chromosome was isolated in the same standard genomic background (from ST8 strain) and re-sequenced in: the second batch. DNA was extracted from 50 adults J1 malesflies using the Nucleobond AXG20 column from Macherey-Nagel. For whole genome sequencing, libraries were prepared using Illumina PCRfree kit and sequencing was done using Illumina HiSeq3000 technology with pair-end 150bp reads. The sequencing depth was determined in order to obtain a minimum autosomal coverage of 80x.

### Mapping and Variant calling

Paired-end reads were trimmed using trimgalore (*trim_galore --paired --nextera --length 75 -- phred33 --no_report_file –fastqc*)(Krueger et al. 2021). Read quality was assessed using FASTQC.

Reads were mapped against the reference genome with bwa (v7.4) using the *BWA-MEM* algorithm (default parameters). We used the heterochromatin-enriched assembly of *D. simulans*(Chang et al. 2022) that include 15Mb of the Y chromosome as reference genome. Resulting sam alignment files were converted into bam files and sorted using respectively samtools (v1.11) *view* and *sort* command. PCR duplicates were removed using *Markduplicates* from Picardtools (v2.12.0) (https://broadinstitute.github.io/picard/). For our SNP calling analyses we want to keep only the uniquely mapping reads, whereas for the repeat quantification we need to keep also the multi-mapping reads. We thus performed two different filtering based on mapping quality using samtools*view*(Li et al. 2009).To keep multi-mapping reads we use the following parameters *-b -h -f 3 -F 4 -F 8 -F 256 -F 2048*, to keep only the uniquely mapping reads we use the following parameters *-b -h -f 3 -F 4 -F 8 -F 256 -F 2048 -q30.* Mean coverage was determined using the *genomecov* tool of bedtools(Quinlan et Hall 2010). The pipeline used (Mapping.sh) can be found on the github repository.

Variant calling was done on the uniquely mapping bam files using *mpileup*and *call* commands from bcftools(v1.14)(Li 2011). We then performed different filtering steps. First isolate the SNPs associated to the Y chromosome using *vcftools*. We use vcftools(Danecek et al. 2011) to analyze the quality of the SNP calling on the Y chromosome based on variant mean depth, variant quality, proportion of missing data per site, mean depth per individual, missing data per individual. This allowed us to determine the threshold for the quality filtering : --remove-indels --max-missing 0.9 --minQ 30 --min-meanDP 10 --max-meanDP 50 --minDP 10 --maxDP 50. Because each iso-Y line originates from a single male and the Y chromosome is supposed to be haploid, heterozygote sites are likely to be duplicated sequences and thus difficult to take into account. We removed the heterozygotes sites using the *filter* command (-e ‘GT=“het”’ option) of bcftools. Note that because Tun83 and Sey81 were sequenced twice, after verification that they were identical, we keep only the data from batch2 to avoid redundancy in the analyses. The pipeline used (VariantCalling.sh) can be found on the github repository.

We calculate the nucleotide diversity along Y chromosomes per 10kb windows using the vcftool*--window-pi 10000* option.

To investigate the population structure we performed a Principal Component Analysis (PCA) on the filtered vcf, using the software plink (v1.9)(Chang et al. 2015). The pairwise distance between strains was calculated using MEGA(Tamura, Stecher, et Kumar 2021).

All the following analyses on the vcf file were done using R scripts (VariantAnalyses.R on github) using the following libraries: *vcfR, SNPrelate, gdsfmt, adegenet, ape, pheatmap, maSigPro, stringr*.

#### Identification of corresponding SNPs in the sister species and phylogeny

To identify the corresponding allele in *D. melanogaster*, *D. sechellia* and *D. mauritiana* we first needed to identify the corresponding location. To do so, we first created a consensus Y chromosome where SNPs are coding in UPAC code, using bcftools*consensus* command (with --iupac-codes option). We built a bed file containing the position of each SNP. Using bedtools*getfasta* command we extracted 400bp sequences for each SNPs (200bp on each side of the SNPs) in a fasta file. Then each sequence were blasted against the Y chromosome consensus of *D. melanogaster*, *D. sechellia* and *D. mauritiana* using blastn (-evalue 1e-20 - qcov_hsp_perc 80 -perc_identity 85 -max_hsps1 -max_target_seqs 1). We output the blast result in a tab delimited format (-outfmt “6 qseqidsseqidevalue length pidentqstartqseqsseq”) and extract the corresponding allele using a R script.

To reconstruct the phylogenetic tree, we first created a fasta sequence with only the SNP for each individual. The SNP for which we were not able to identify a corresponding allele in the sister species were coded as gaps. We constructed the maximum likelihood tree using the GTR model and ultrafast bootstrap (1000 replicates) using iqTREE web server(Nguyen et al. 2015).

*D. melanogaster* was used as the outgroup. The scripts used can be found in github : Figure3/Ancestry.sh and Figure3/Ancestry.R.

#### Transposable element and complex satellite quantification

Due to the batch effect observed we only used the strain from the second batch for the analysis. To estimate the abundance of transposable elements and complex satellites in each iso-Y line we first mapped the read against our reference genome. We applied the same pipeline as for the SNP calling, with the exception that we kept multi-mapping reads (pipeline Mappingt.sh on github). The reference genome was annotated using a custom repeat library (SupFile1.fasta) using repeatmasker (options *-no_is -a -inv -pa 20 -div 20*)(Smit, AFA, Hubley, R, et Green, P. 2013). Using htseq-count (Putri et al. 2022)we counted the number of reads that map against each repeats and normalize the reads count in RPM. We removed repeats for none of the strain have a RPM greater than 10. All the subsequent analyses were done using Rstudio. We performed a Wilcoxon test, followed by an “fdr” correction, to determine which repeats have a different enrichment between sensitive and resistant. We calculated the coefficient of variation for each repeat to determine which element should be the most variable, the top 25 is presented in Figure 4D. The scripts used can be found in github : TEcount.sh and Figure4D/TE.R.

#### kmers quantification

Due to the batch effect observed we only used the strain from the second batch for the analysis. Barring the fourth chromosome (not controlled), only the Y chromosome differs; we thus consider that variations observed between lines are due to the Y chromosome. To estimate the abundance of simple satellites in each strain we used an assembly-free approach based on kmer quantification. We used the software k-Seek which quantified the simple satellite tandemly repeated using unassembled Illumina reads. We normalized the kmers count in using the total mapping reads. We removed the kmers for which none of the strain have anormalized count greater than 5. All the subsequent analyses were done using Rstudio. We performed a Wilcoxon test to determine which kmers have a different enrichment between sensitive and resistant. Then we performed a correlation test between the kmers count and the resistance ability for the 11 strains that have been phenotype (Figure 1B, Supplementary Table1). The scripts used can be found in github : kmers.sh and Figure5/Kmers.R.

#### Gene copy number analysis

Due to the batch effect observed we only used the strain from the second batch for the analysis. To estimate the exon copy number in each iso-Y line we first mapped the reads against our reference genome. We applied the same pipeline as for the SNP calling, with the exception that we kept multi-mapping reads (pipeline Mappingt.sh on github). We estimated the reads coverage on each exon using *samtools depth*. For the 11 canonical genes we used a bed file containing the coordinate of each exon on the reference genome based on Chang et al 2022. For the two ampliconic genes, we blasted the consensus sequences against the reference genome (option *-perc_identity 80*) to determine the coordinate of each insertion. For each iso-Y line we calculated the mean coverage of all single copy exons and used it as a normalization factor. To estimate the exon copy number we calculated the exon mean coverage divided by the normalization factor and multiplied by the number of corresponding exons annotated on the reference genome. The script used can be found in github : Figure4A/Gene_count.R.

#### Genotyping and Haplotype tree

We designed primers around SNPs located on regions annotated as coding regions (Chang et al 2022), as they are the most likely to be unique. For each of them we performed a PCR with DreamTaq™ Hot Start Green PCR (ThermoFisherScientific, ref K9021) following manufacturer’s instructions (hybridization temperature: 58°). Each PCR product was sequenced using the Sanger method on a SeqStudio Genetic Analyzer (ThermoScientific). Only five primers pairs were able to deliver a unique sequence (Supplementary Table6), containing 11 SNPs in total (Supplementary Table2). We genotyped the sequenced iso-Y lines, to validate our SNP calling, and 86 additional iso-Y lines from 22 populations. Similarly, we genotyped the mitochondrial haplotype of the 21 iso-Y lines using 2 pairs of primers in Supplementary Table6. We concatenated the 11 SNPs for each iso-Y strains in a nexus format and built the haplotype tree by the Median Joining Method using POPart(Leigh et Bryant 2015).

#### Fluorescence in situ hybridization

The FISH was performed using primary probe for (AATGG)n and (AAACAAT)n coupled with sec6 and sec5 adaptors. The sequences for each probe are in Supplementary Table7. Sec5 is coupled with Cy5 while sec6 is coupled with Cy3. We dissected brains from third instar larvae in PBS, incubate 8min in 0.5% Sodium citrate. We fix for 6 min in 4% formaldehyde, 45% acetic acid before squashing. The brains were put between the slide and coverslip and squashed hard before being immersed in liquid nitrogen. After 10 min in 100% ethanol, slides were air dried for at least one hour before proceeding to the hybridization. For the hybridization we use 20pmol of primary probes and 80pmol of the secondary probes in 50 ul of hybridization buffer (50% formamide, 10% dextran sulfate, 2xSSC). Slides were heated for 5 min at 95°C for denaturation and incubated overnight at 37°C in humid chamber. Slides were then washed 3 times 5min with 4XSSCT and 3 times 5min with 0.1SSC before being mounted in slowfade DAPI. We imaged using LEICA DM5500 microscope and edited the image using Fiji.

#### Slot blot

We performed the slot blot for 14 iso-Y strains. We extracted the DNA from 30 adult males using the QIAGEN DNeasy tissue and blood kit. We use 200ng of DNA for the slot blot. First the DNA was incubated 10 min in denaturing buffer (0.4 NaOH, 25 mM EDTA) at room temperature then cooled down in ice cold loading buffer (0.1X SSC, 0.125 N NaOH). The blot was performed using the slot blotter apparatus from BioRad. The membrane was UV cross-linked for 3 min. Hybridization was performed overnight at 42°C using 15 ml of hybridization buffer (ULTRAhyb, Thermo Scientific) and 100pmol of primary probes and 200pmol of secondary fluorescent probes (Supplementary Table7). The next day we washed the membrane 3 times 15 min using the wash buffer (North2South™ Hybridization Stringency Wash Buffer, Thermo Scientific). We imaged the membrane using Odyssey XL. For each strain, we made 3 replicates that were randomly placed on the blot. The intensity of the simple satellite signals were normalized by the intensity of our control gene, *rp49*.

## Data availability

All sequences are available from NCBI SRA under Bioproject accession PRJNA905841. All the BASH pipelines and R scripts used in this study are available on github https://github.com/CourretC/YchromosomesCourret2023.git

## Acknowledgment

This research has been supported by the Centre National de la Recherche Scientifique (UMR9191) and by grants from the AgenceNationale de la Recherche (RESIST ANR-21-CE02-0004-02) and by National Institutes of HealthNIH R35GM119515to AML.

## SUPPLEMENTAL FIGURES

<u>Supplementary Figure 1:</u> Y chromosome haplotypes. Each line corresponds to a different Y chromosome ordered based on a clustering method. Each column corresponds to a SNP, the order of which is based on their physical position on the Y chromosome. The blue alleles correspond to the reference allele and the red alleles correspond to the alternative allele; white are missing data.

<u>Supplementary Figure 2 :</u>Haplotype network of 98 Y chromosomes from 26 populations. The haplotypes are composed of 11 SNPs. The name of each Y chromosome is indicated within each cluster (see Supplementary Table2 for full name). The clusters are colored according to the 4 clusters identified for the 21 iso-Y lines sequenced.

<u>Supplementary Figure 3:</u> The box plots represent the normalized abundance of satellite abundance (y axis) in each group (resistant Y’s in red and sensitive Y’s in blue).

<u>Supplementary Figure 4 :</u> Correlation plots between the satellite abundance (y axis) and the resistance ability (x-axis). Correlation was calculated based on a Spearman correlation test

<u>Supplementary Figure 5 :</u> FISH on mitotic chromosomes from larval brains using (AACAAT)_4_ (green) and (AATGG)_6_ (red) probes.

<u>Supplementary Figure 6 :</u> FISH on mitotic chromosomes from larval brains using (AAGAGAAGAGAG)_4_ (green) and (AACAATT)_64_ (red) probes.

<u>Supplementary Figure 7 :</u> FISH on mitotic chromosomes from larval brains using (AACAATC)_4_ (green) and (AAACAAC)_4_ (red) probes.

<u>Supplementary Figure 8:</u> Slot blot with DNA from 14 iso-Y strains, 3 replicates for each strain. The y-axis correspond the signal from the hybridization using probes targeting AAACAAC (A), AAACAAT (B) and AACAATC (C). The error bar correspond to the standard error. The signal intensity was normalized by the rp49 signal and plotted against the relative abundance (in log10) inferred by k-seek. The correlation was calculated based on Pearson correlation test, the red line corresponds to the regression line.

## Reference

1. Agudo, M., A. Losada, J. P. Abad, S. Pimpinelli, P. Ripoll, et A. Villasante. 1999. « Centromeres from Telomeres? The Centromeric Region of the Y Chromosome of Drosophila Melanogaster Contains a Tandem Array of Telomeric HeT-A- and TART-Related Sequences ». Nucleic Acids Research 27 (16): 3318–24. https://doi.org/10.1093/nar/27.16.3318.

2. Bachtrog, Doris. 2013. « Y-Chromosome Evolution: Emerging Insights into Processes of Y-Chromosome Degeneration ». Nature Reviews Genetics 14 (2): 113–24. https://doi.org/10.1038/nrg3366.

3. Bastide, Héloïse, Michel Cazemajor, David Ogereau, Nicolas Derome, Frédéric Hospital, et Catherine Montchamp-Moreau. 2011. « Rapid Rise and Fall of Selfish Sex-Ratio X Chromosomes in Drosophila Simulans: Spatiotemporal Analysis of Phenotypic and Molecular Data ». Molecular Biology and Evolution 28 (9): 2461–70. https://doi.org/10.1093/molbev/msr074.

4. Bastide, Héloïse, Pierre R. Gérard, David Ogereau, Michel Cazemajor, et Catherine Montchamp-Moreau. 2013. « Local Dynamics of a Fast-Evolving *Sex-Ratio* System in *Drosophila Simulans* ». Molecular Ecology 22 (21): 5352–67. https://doi.org/10.1111/mec.12492.

5. Bernardo Carvalho, A., Leonardo B. Koerich, et Andrew G. Clark. 2009. « Origin and Evolution of Y Chromosomes: Drosophila Tales ». Trends in Genetics 25 (6): 270–77. https://doi.org/10.1016/j.tig.2009.04.002.

6. Boulétreau-Merle, Josselyne, Pierre Fouillet, et Julien Varaldi. 2003. « Divergent Strategies in Low Temperature Environment for the Sibling Species Drosophila Melanogaster and D. Simulans: Overwintering in Extension Border Areas of France and Comparison with African Populations ». Evolutionary Ecology 17 (5–6): 523–48. https://doi.org/10.1023/B:EVEC.0000005632.21186.21.

7. Branco, A T, Y Tao, D L Hartl, et B Lemos. 2013. « Natural Variation of the Y Chromosome Suppresses Sex Ratio Distortion and Modulates Testis-Specific Gene Expression in Drosophila Simulans ». Heredity 111 (1): 8–15. https://doi.org/10.1038/hdy.2013.5.

8. Branco, Alan T., Rute M Brito, et Bernardo Lemos. 2017. « Sex-Specific Adaptation and Genomic Responses to Y Chromosome Presence in Female Reproductive and Neural Tissues ». Proceedings of the Royal Society B: Biological Sciences 284 (1869): 20172062. https://doi.org/10.1098/rspb.2017.2062.

9. Broad Institute. 2019. « Picard Toolkit », 2019, Broad Institute, GitHub repository édition. https://broadinstitute.github.io/picard/.

10. Brown, Emily J, Alison H Nguyen, et Doris Bachtrog. 2020. « The Drosophila Y Chromosome Affects Heterochromatin Integrity Genome-Wide ». Édité par John Parsch. Molecular Biology and Evolution 37 (10): 2808–24. https://doi.org/10.1093/molbev/msaa082.

11. Cazemajor, Michel, Claudie Landré, et Catherine Montchamp-Moreau. 1997. « The *Sex-Ratio* Trait in *Drosophila Simulans:* Genetic Analysis of Distortion and Suppression ». Genetics 147 (2): 635–42. https://doi.org/10.1093/genetics/147.2.635.

12. Chang, Ching-Ho, Lauren E Gregory, Kathleen E Gordon, Colin D Meiklejohn, et Amanda M Larracuente. 2022. « Unique Structure and Positive Selection Promote the Rapid Divergence of Drosophila Y Chromosomes ». ELife 11 (janvier): e75795. https://doi.org/10.7554/eLife.75795.

13. Chang, Ching-Ho, et Amanda M. Larracuente. 2019. « Heterochromatin-Enriched Assemblies Reveal the Sequence and Organization of the *Drosophila Melanogaster* Y Chromosome ». Genetics 211 (1): 333–48. https://doi.org/10.1534/genetics.118.301765.

14. Chang, Christopher C, Carson C Chow, Laurent CAM Tellier, Shashaank Vattikuti, Shaun M Purcell, et James J Lee. 2015. « Second-Generation PLINK: Rising to the Challenge of Larger and Richer Datasets ». GigaScience 4 (1): 7. https://doi.org/10.1186/s13742-015-0047-8.

15. Charlesworth, B., J. A. Coyne, et N. H. Barton. 1987. « The Relative Rates of Evolution of Sex Chromosomes and Autosomes ». The American Naturalist 130 (1): 113–46. https://doi.org/10.1086/284701.

16. Clark, A G. 1990. « Two Tests of Y Chromosomal Variation in Male Fertility of Drosophila Melanogaster. » Genetics 125 (3): 527–34. https://doi.org/10.1093/genetics/125.3.527.

17. Clark, Andrew G. 1987. « Variation in *Y* Chromosome Segregation in Natural Populations of *Drosophila Melanogaster* ». Genetics 115 (1): 143–51. https://doi.org/10.1093/genetics/115.1.143.

18. Courret, Cécile, Ching-Ho Chang, Kevin H.-C. Wei, Catherine Montchamp-Moreau, et Amanda M. Larracuente. 2019. « Meiotic Drive Mechanisms: Lessons from *Drosophila* ». Proceedings of the Royal Society B: Biological Sciences 286 (1913): 20191430. https://doi.org/10.1098/rspb.2019.1430.

19. Danecek, P., A. Auton, G. Abecasis, C. A. Albers, E. Banks, M. A. DePristo, R. E. Handsaker, et al. 2011. « The Variant Call Format and VCFtools ». Bioinformatics 27 (15): 2156–58. https://doi.org/10.1093/bioinformatics/btr330.

20. Dean, Matthew D., et J. William O. Ballard. 2004. « Linking Phylogenetics with Population Genetics to Reconstruct the Geographic Origin of a Species ». Molecular Phylogenetics and Evolution 32 (3): 998–1009. https://doi.org/10.1016/j.ympev.2004.03.013.

21. Fouvry, Lucie, David Ogereau, Anne Berger, Frederick Gavory, et Catherine Montchamp-Moreau. 2011. « Sequence Analysis of the Segmental Duplication Responsible for Paris *Sex-Ratio* Drive in *Drosophila Simulans* ». G3 Genes|Genomes|Genetics 1 (5): 401–10. https://doi.org/10.1534/g3.111.000315.

22. Francisco, Flávio O., et Bernardo Lemos. 2014. « How Do Y-Chromosomes Modulate Genome-Wide Epigenetic States: Genome Folding, Chromatin Sinks, and Gene Expression ». Journal of Genomics 2: 94–103. https://doi.org/10.7150/jgen.8043.

23. Hall, David W. 2004. « MEIOTIC DRIVE AND SEX CHROMOSOME CYCLING ». Evolution 58 (5): 925–31. https://doi.org/10.1111/j.0014-3820.2004.tb00426.x.

24. Hamblin, Martha T, et Michel Veuille. 1999. « Population Structure Among African and Derived Populations of Drosophila Simulans: Evidence for Ancient Subdivision and Recent Admixture ». Genetics 153 (1): 305–17. https://doi.org/10.1093/genetics/153.1.305.

25. Hartl, Daniel L. 1974. « GENETIC DISSECTION OF SEGREGATION DISTORTION. I. SUICIDE COMBINATIONS OF SD GENES ». Genetics 76 (3): 477–86. https://doi.org/10.1093/genetics/76.3.477.

26. Helleu, Quentin, Cécile Courret, David Ogereau, Katie L Burnham, Nicole Chaminade, Mohamed Chakir, Sylvie Aulard, et Catherine Montchamp-Moreau. 2019. « Sex-Ratio Meiotic Drive Shapes the Evolution of the Y Chromosome in Drosophila Simulans ». Molecular Biology and Evolution 36 (12): 2668–81. https://doi.org/10.1093/molbev/msz160.

27. Helleu, Quentin, Pierre R. Gérard, Raphaëlle Dubruille, David Ogereau, Benjamin Prud’homme, Benjamin Loppin, et Catherine Montchamp-Moreau. 2016. « Rapid Evolution of a Y-Chromosome Heterochromatin Protein Underlies Sex Chromosome Meiotic Drive ». Proceedings of the National Academy of Sciences 113 (15): 4110–15. https://doi.org/10.1073/pnas.1519332113.

28. Helleu, Quentin, Pierre R. Gérard, et Catherine Montchamp-Moreau. 2015. « Sex Chromosome Drive ». Cold Spring Harbor Perspectives in Biology 7 (2): a017616. https://doi.org/10.1101/cshperspect.a017616.

29. Hill, Tom, Christian Schlötterer, et Andrea J. Betancourt. 2016. « Hybrid Dysgenesis in Drosophila Simulans Associated with a Rapid Invasion of the P-Element ». Édité par Harmit S. Malik. PLOS Genetics 12 (3): e1005920. https://doi.org/10.1371/journal.pgen.1005920.

30. Hoskins, Roger A, Christopher D Smith, Joseph W Carlson, A Bernardo Carvalho, Aaron Halpern, Joshua S Kaminker, Cameron Kennedy, et al. 2002. « Heterochromatic sequences in a Drosophila whole-genome shotgun assembly. » Genome Biology 3 (12): research0085.1. https://doi.org/10.1186/gb-2002-3-12-research0085.

31. Huttunen, Susanna, et Jouni Aspi. s. d. « Complex Inheritance of Male Courtship Song Characters in Drosophila Virilis », 8.

32. Irvin, Steven D, Kris A Wetterstrand, Carolyn M Hutter, et Charles F Aquadro. 1998. « Genetic Variation and Differentiation at Microsatellite Loci in Drosophila Simulans: Evidence for Founder Effects in New World Populations ». Genetics 150 (2): 777–90. https://doi.org/10.1093/genetics/150.2.777.

33. Kennison, James A. 1981. « THE GENETIC AND CYTOLOGICAL ORGANIZATION OF THE *Y* CHROMOSOME OF *DROSOPHILA MELANOGASTER* ». Genetics 98 (3): 529–48. https://doi.org/10.1093/genetics/98.3.529.

34. Kopp, Artyom, Amanda Frank, et Jeffrey Fu. 2006. « Historical Biogeography of Drosophila Simulans Based on Y-Chromosomal Sequences ». Molecular Phylogenetics and Evolution 38 (2): 355–62. https://doi.org/10.1016/j.ympev.2005.06.006.

35. Krueger, Felix, Frankie James, Phil Ewels, Ebrahim Afyounian, et Benjamin Schuster-Boeckler. 2021. « FelixKrueger/TrimGalore: v0.6.7 - DOI via Zenodo ». Zenodo. https://doi.org/10.5281/ZENODO.5127899.

36. Lachaise, Daniel, et Jean-François Silvain. s. d. « How Two Afrotropical Endemics Made Two Cosmopolitan Human Commensals: The Drosophila Melanogaster–D. Simulans Palaeogeographic Riddle », 23.

37. Larracuente, Amanda M, et Andrew G Clark. 2013. « Surprising Differences in the Variability of Y Chromosomes in African and Cosmopolitan Populations of *Drosophila Melanogaster* ». Genetics 193 (1): 201–14. https://doi.org/10.1534/genetics.112.146167.

38. Leigh, Jessica W., et David Bryant. 2015. « POPART : Full-feature Software for Haplotype Network Construction ». Édité par Shinichi Nakagawa. Methods in Ecology and Evolution 6 (9): 1110–16. https://doi.org/10.1111/2041-210X.12410.

39. Lemos, Bernardo, Alan T Branco, Pan-Pan Jiang, Daniel L Hartl, et Colin D Meiklejohn. 2014. « Genome-Wide Gene Expression Effects of Sex Chromosome Imprinting in Drosophila ». G3 Genes|Genomes|Genetics 4 (1): 1–10. https://doi.org/10.1534/g3.113.008029.

40. Li, H. 2011. « A Statistical Framework for SNP Calling, Mutation Discovery, Association Mapping and Population Genetical Parameter Estimation from Sequencing Data ». Bioinformatics 27 (21): 2987–93. https://doi.org/10.1093/bioinformatics/btr509.

41. Li, H., B. Handsaker, A. Wysoker, T. Fennell, J. Ruan, N. Homer, G. Marth, G. Abecasis, R. Durbin, et 1000 Genome Project Data Processing Subgroup. 2009. « The Sequence Alignment/Map Format and SAMtools ». Bioinformatics 25 (16): 2078–79. https://doi.org/10.1093/bioinformatics/btp352.

42. Lima, Leonardo G de, Stacey L Hanlon, et Jennifer L Gerton. 2020. « Origins and Evolutionary Patterns of the 1.688 Satellite DNA Family in Drosophila Phylogeny ». G3 Genes|Genomes|Genetics 10 (11): 4129–46. https://doi.org/10.1534/g3.120.401727.

43. Lyttle, T W. 1989. « The Effect of Novel Chromosome Position and Variable Dose on the Genetic Behavior of the Responder (Rsp) Element of the Segregation Distorter (SD) System of Drosophila Melanogaster. » Genetics 121 (4): 751–63. https://doi.org/10.1093/genetics/121.4.751.

44. Machado, Heather E., Alan O. Bergland, Katherine R. O’Brien, Emily L. Behrman, Paul S. Schmidt, et Dmitri A. Petrov. 2016. « Comparative Population Genomics of Latitudinal Variation in Drosophila Simulans and Drosophila Melanogaster ». Molecular Ecology 25 (3): 723–40. https://doi.org/10.1111/mec.13446.

45. Meiklejohn, Colin D, Emily L Landeen, Kathleen E Gordon, Thomas Rzatkiewicz, Sarah B Kingan, Anthony J Geneva, Jeffrey P Vedanayagam, et al. 2018. « Gene Flow Mediates the Role of Sex Chromosome Meiotic Drive during Complex Speciation ». ELife 7 (décembre): e35468. https://doi.org/10.7554/eLife.35468.

46. Montchamp-Moreau, Catherine, et Michel Cazemajor. 2002. « *Sex-Ratio* Drive in *Drosophila Simulans* : Variation in Segregation Ratio of *X* Chromosomes From a Natural Population ». Genetics 162 (3): 1221–31. https://doi.org/10.1093/genetics/162.3.1221.

47. Montchamp-Moreau, Catherine, Valérie Ginhoux, et Anne Atlan. 2001. « THE Y CHROMOSOMES OF DROSOPHILA SIMULANS ARE HIGHLY POLYMORPHIC FOR THEIR ABILITY TO SUPPRESS SEX-RATIO DRIVE ». Evolution 55 (4): 728. https://doi.org/10.1554/0014-3820(2001)055[0728:TYCODS]2.0.CO;2.

48. Muirhead, Christina A., et Daven C. Presgraves. 2021. « Satellite DNA-Mediated Diversification of a Sex-Ratio Meiotic Drive Gene Family in Drosophila ». Nature Ecology & Evolution 5 (12): 1604–12. https://doi.org/10.1038/s41559-021-01543-8.

49. Nguyen, Lam-Tung, Heiko A. Schmidt, Arndt von Haeseler, et Bui Quang Minh. 2015. « IQ-TREE: A Fast and Effective Stochastic Algorithm for Estimating Maximum-Likelihood Phylogenies ». Molecular Biology and Evolution 32 (1): 268–74. https://doi.org/10.1093/molbev/msu300.

50. Putri, Givanna H, Simon Anders, Paul Theodor Pyl, John E Pimanda, et Fabio Zanini. 2022. « Analysing High-Throughput Sequencing Data in Python with HTSeq 2.0 ». Édité par Valentina Boeva. Bioinformatics 38 (10): 2943–45. https://doi.org/10.1093/bioinformatics/btac166.

51. Quinlan, Aaron R., et Ira M. Hall. 2010. « BEDTools: A Flexible Suite of Utilities for Comparing Genomic Features ». Bioinformatics 26 (6): 841–42. https://doi.org/10.1093/bioinformatics/btq033.

52. R Core Team. 2022. « R: A language and environment for statistical computing. », 2022. https://www.R-project.org/.

53. Rice, William R. 2014. « An X-Linked Sex Ratio Distorter in *Drosophila Simulans* That Kills or Incapacitates Both Noncarrier Sperm and Sons ». G3 Genes|Genomes|Genetics 4 (10): 1837–48. https://doi.org/10.1534/g3.114.013292.

54. Rohmer, Céline, Jean R. David, Brigitte Moreteau, et Dominique Joly. 2004. « Heat Induced Male Sterility in *Drosophila Melanogaster* : Adaptive Genetic Variations among Geographic Populations and Role of the Y Chromosome ». Journal of Experimental Biology 207 (16): 2735–43. https://doi.org/10.1242/jeb.01087.

55. Smit, AFA, Hubley, R, et Green, P. 2013. « RepeatMasker Open-4.0. », 2015 2013. http://www.repeatmasker.org.

56. Solignac, Michel, et Monique Monnerot. 1986. « RACE FORMATION, SPECIATION, AND INTROGRESSION WITHIN *DROSOPHILA SIMULANS, D. MAURITIANA*, AND *D. SECHELLIA* INFERRED FROM MITOCHONDRIAL DNA ANALYSIS ». Evolution 40 (3): 531–39. https://doi.org/10.1111/j.1558-5646.1986.tb00505.x.

57. Sproul, John S, Danielle E Khost, Danna G Eickbush, Sherif Negm, Xiaolu Wei, Isaac Wong, et Amanda M Larracuente. 2020. « Dynamic Evolution of Euchromatic Satellites on the X Chromosome in Drosophila Melanogaster and the Simulans Clade ». Édité par John Parsch. Molecular Biology and Evolution 37 (8): 2241–56. https://doi.org/10.1093/molbev/msaa078.

59. Tamura, Koichiro, Glen Stecher, et Sudhir Kumar. 2021. « MEGA11: Molecular Evolutionary Genetics Analysis Version 11 ». Édité par Fabia Ursula Battistuzzi. Molecular Biology and Evolution 38 (7): 3022–27. https://doi.org/10.1093/molbev/msab120.

60. Tao, Yun, Luciana Araripe, Sarah B Kingan, Yeyan Ke, Hailian Xiao, et Daniel L Hartl. 2007. « A Sex-Ratio Meiotic Drive System in Drosophila Simulans. II: An X-Linked Distorter ». Édité par Daniel Barbash. PLoS Biology 5 (11): e293. https://doi.org/10.1371/journal.pbio.0050293.

61. Vedanayagam, Jeffrey, Ching-Jung Lin, et Eric C. Lai. 2021. « Rapid Evolutionary Dynamics of an Expanding Family of Meiotic Drive Factors and Their HpRNA Suppressors ». Nature Ecology & Evolution 5 (12): 1613–23. https://doi.org/10.1038/s41559-021-01592-z.

62. Wall, Jeffrey D, Peter Andolfatto, et Molly Przeworski. 2002. « Testing Models of Selection and Demography in *Drosophila Simulans* ». Genetics 162 (1): 203–16. https://doi.org/10.1093/genetics/162.1.203.

63. Wei, Kevin H -C, Sarah E Lower, Ian V Caldas, Trevor J S Sless, Daniel A Barbash, et Andrew G Clark. 2018. « Variable Rates of Simple Satellite Gains across the Drosophila Phylogeny ». Molecular Biology and Evolution 35 (4): 925–41. https://doi.org/10.1093/molbev/msy005.

64. Wei, Kevin H.-C., Jennifer K. Grenier, Daniel A. Barbash, et Andrew G. Clark. 2014. « Correlated Variation and Population Differentiation in Satellite DNA Abundance among Lines of *Drosophila Melanogaster* ». Proceedings of the National Academy of Sciences 111 (52): 18793–98. https://doi.org/10.1073/pnas.1421951112.

65. Zurovcova, Martina, et Walter F Eanes. 1999. « Lack of Nucleotide Polymorphism in the Y-Linked Sperm Flagellar Dynein Gene Dhc-Yh3 of Drosophila Melanogaster and D. Simulans ». Genetics 153 (4): 1709–15. https://doi.org/10.1093/genetics/153.4.1709.

