## Supplementary figures and images for "The evolutionary history of *Drosophila simulans* Y chromosomes reveals molecular signatures of resistance to sex ratio meiotic drive"

### Supplemental Figure 1

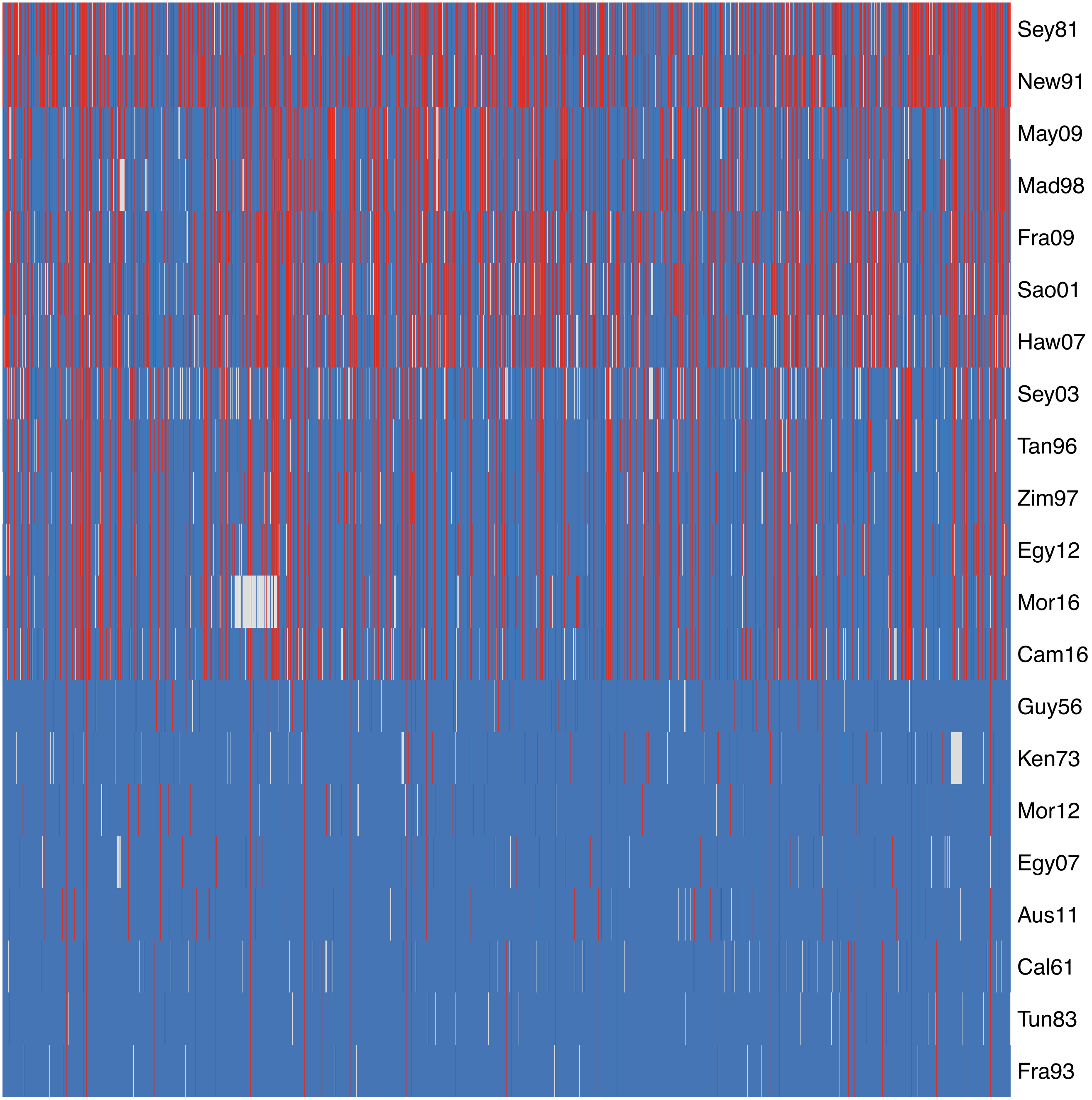

### Supplemental Figure 2

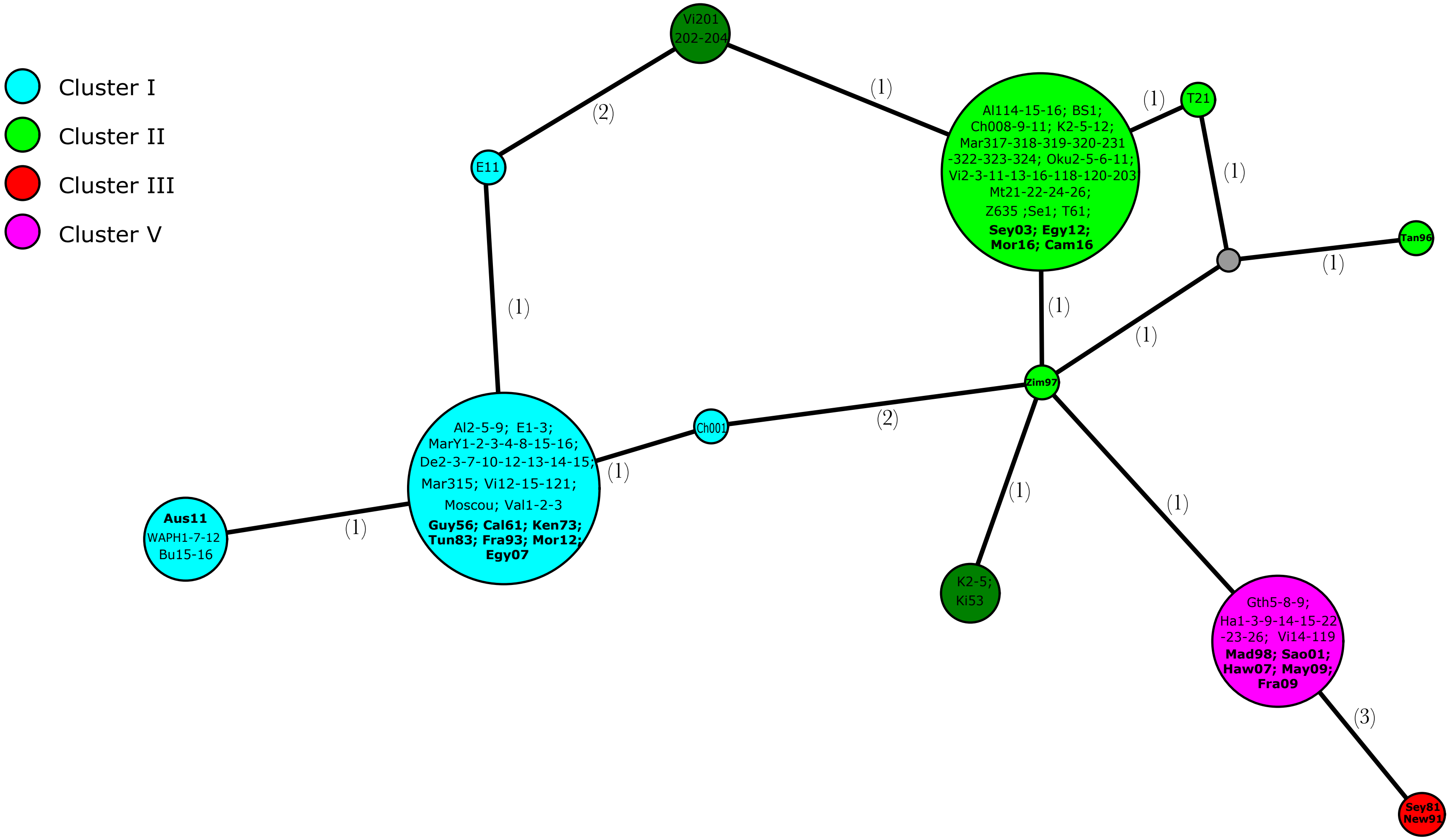

### Supplemental Figure 3

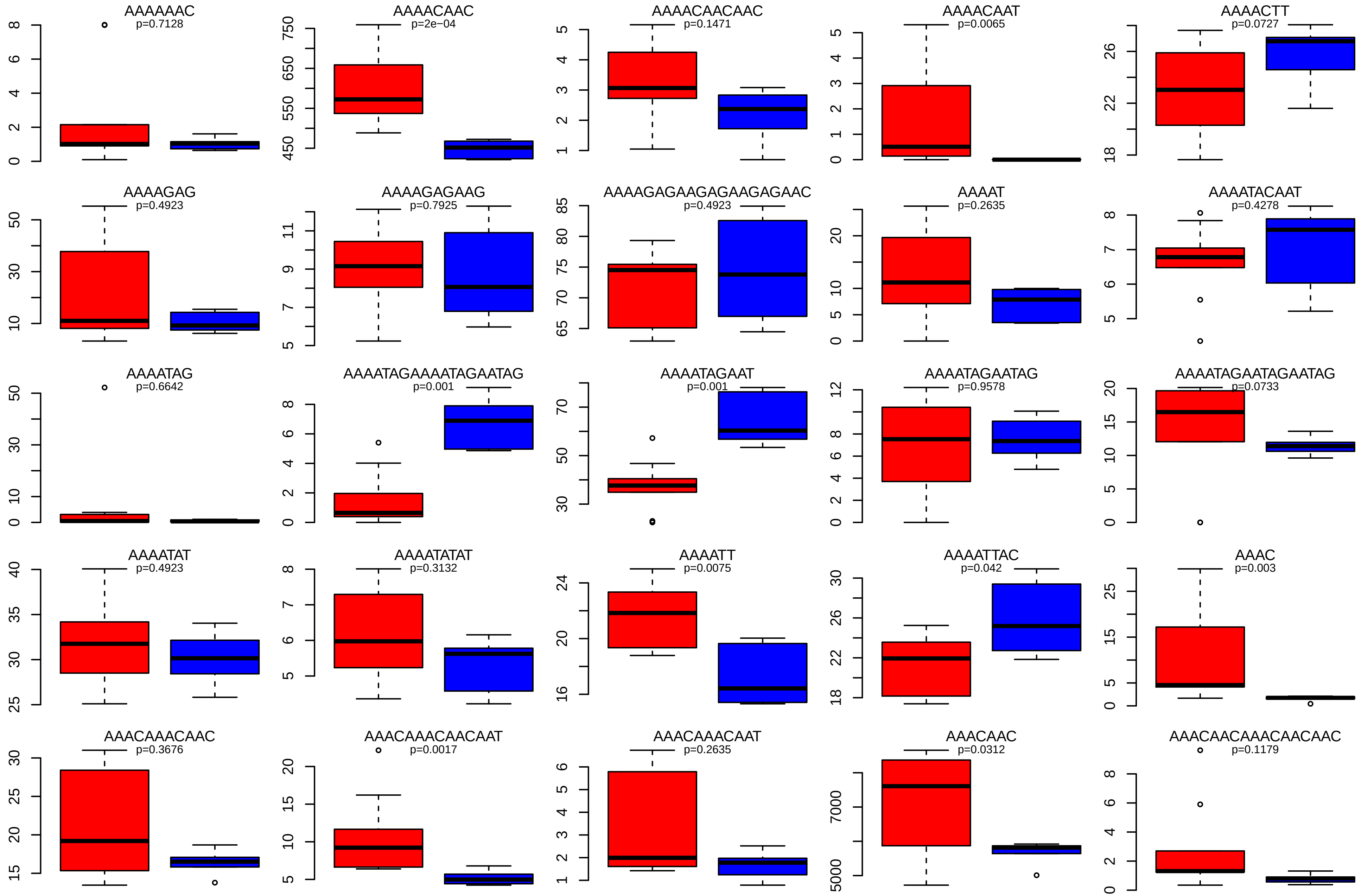

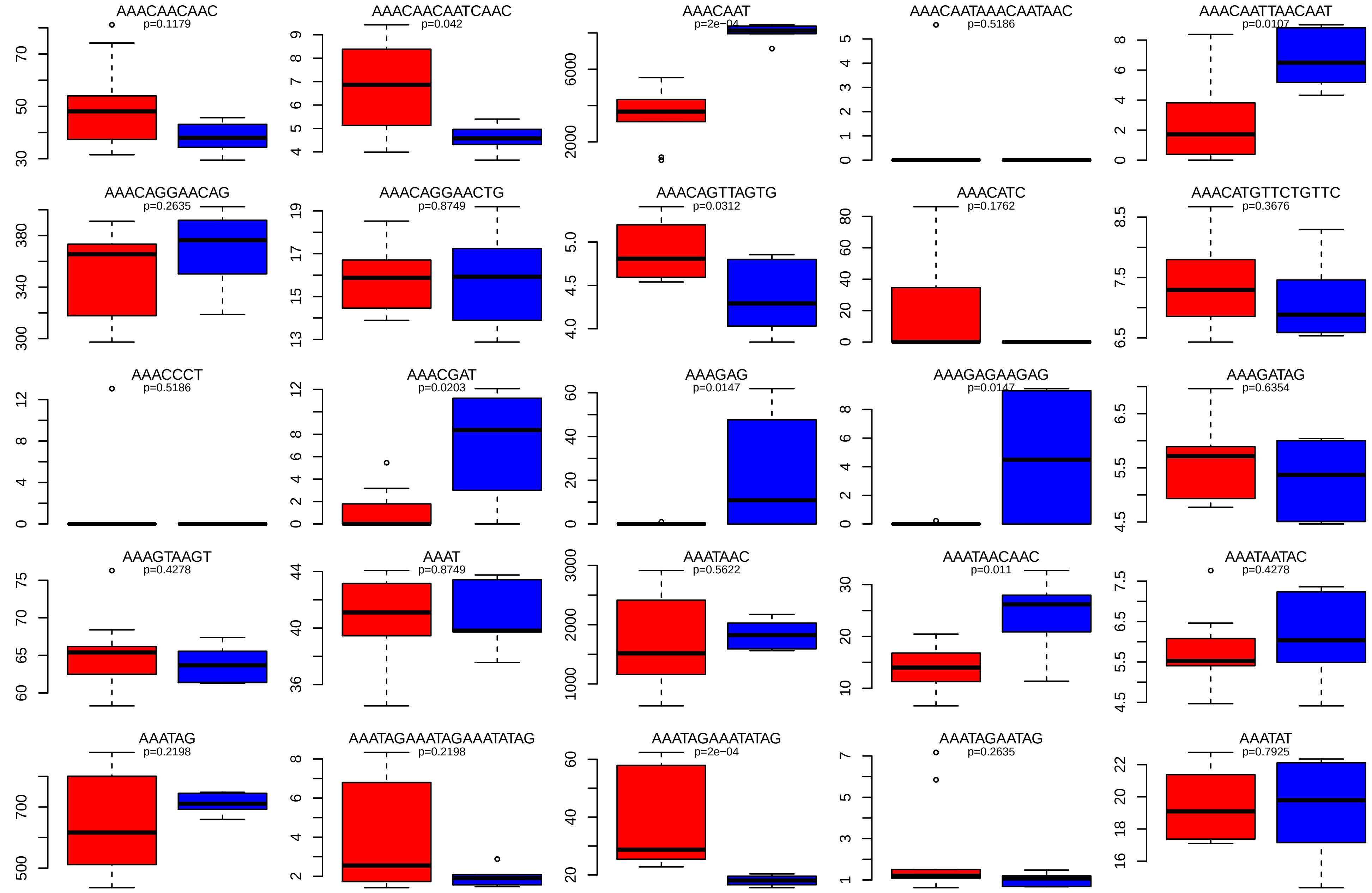

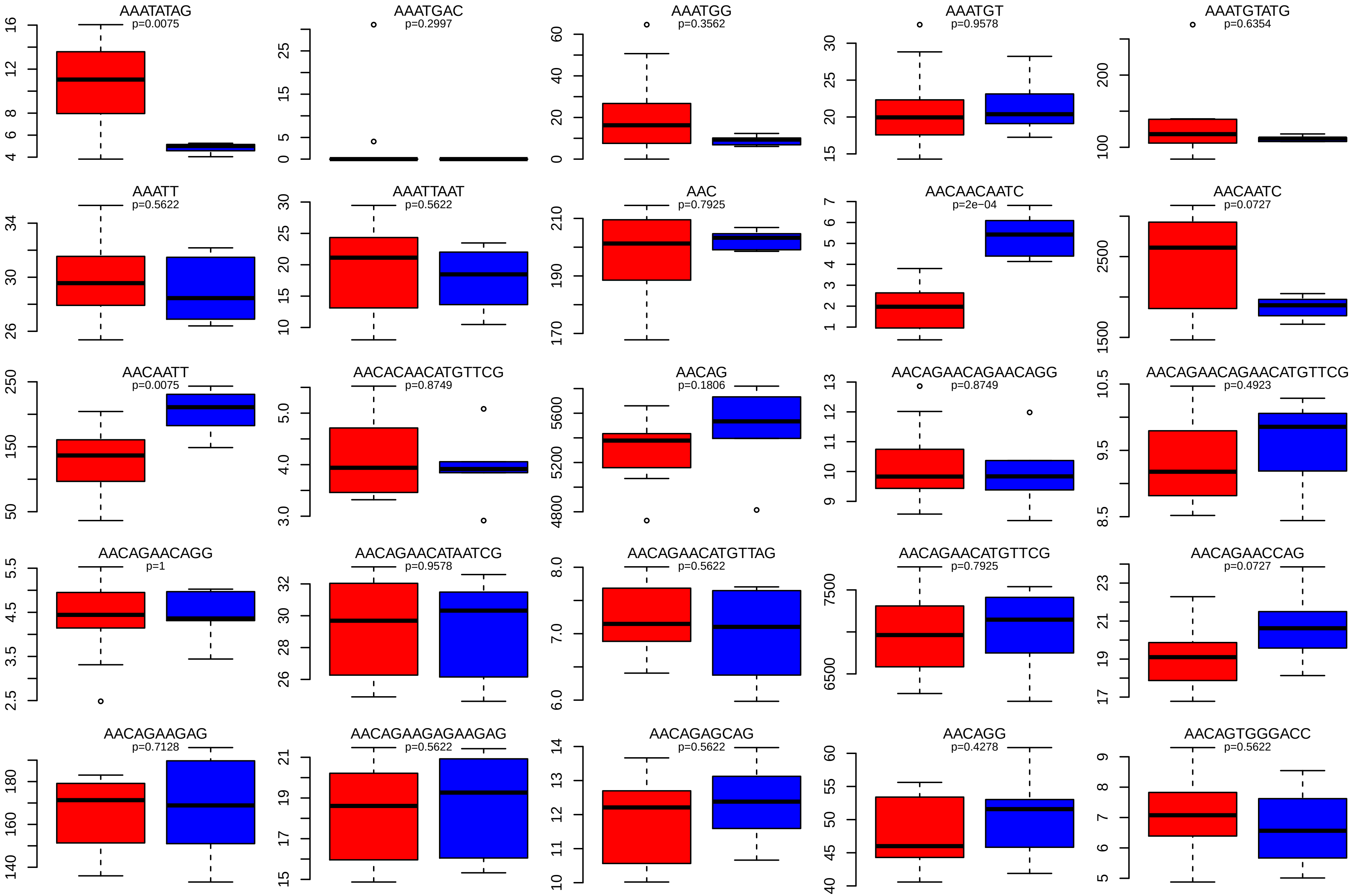

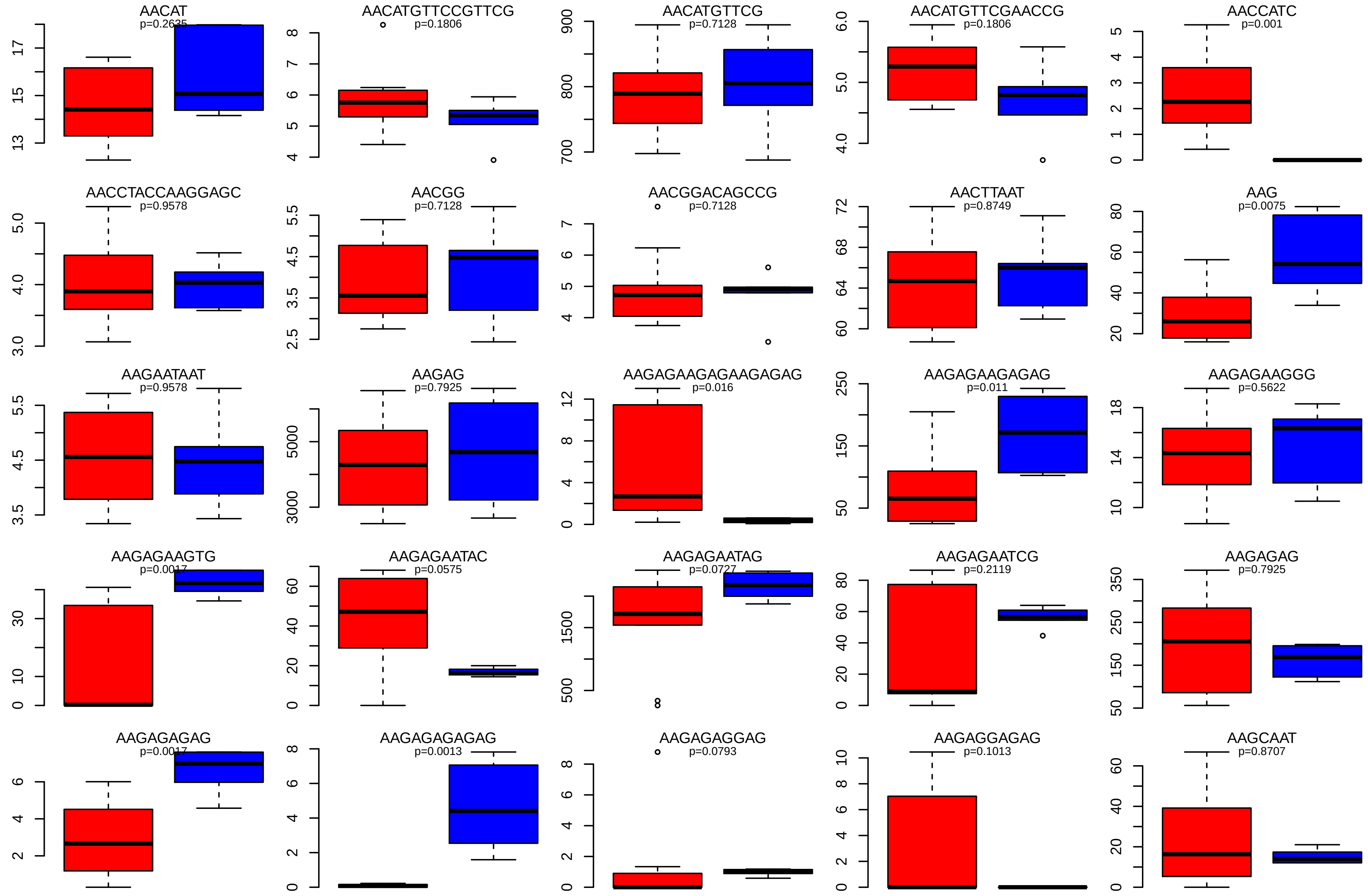

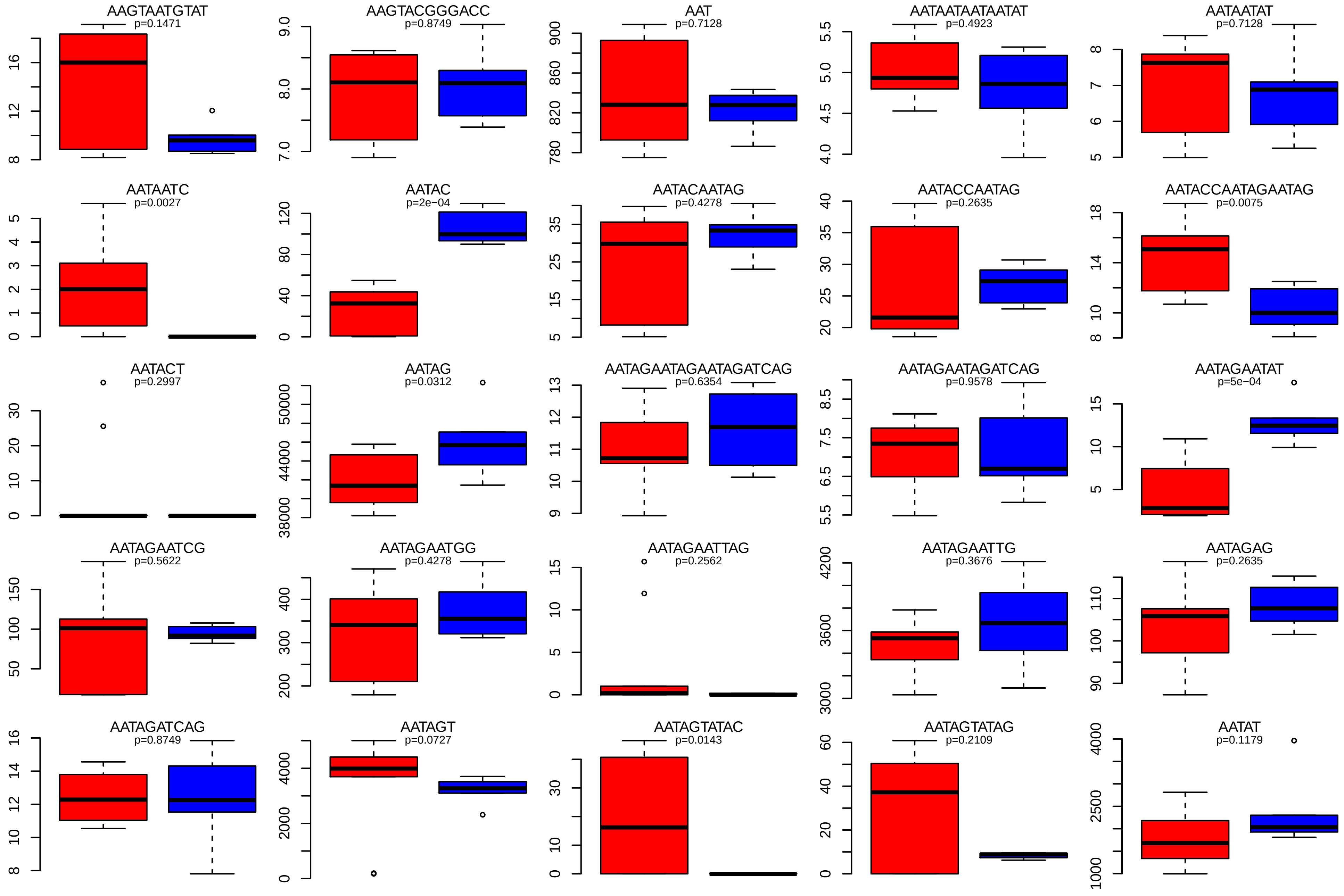

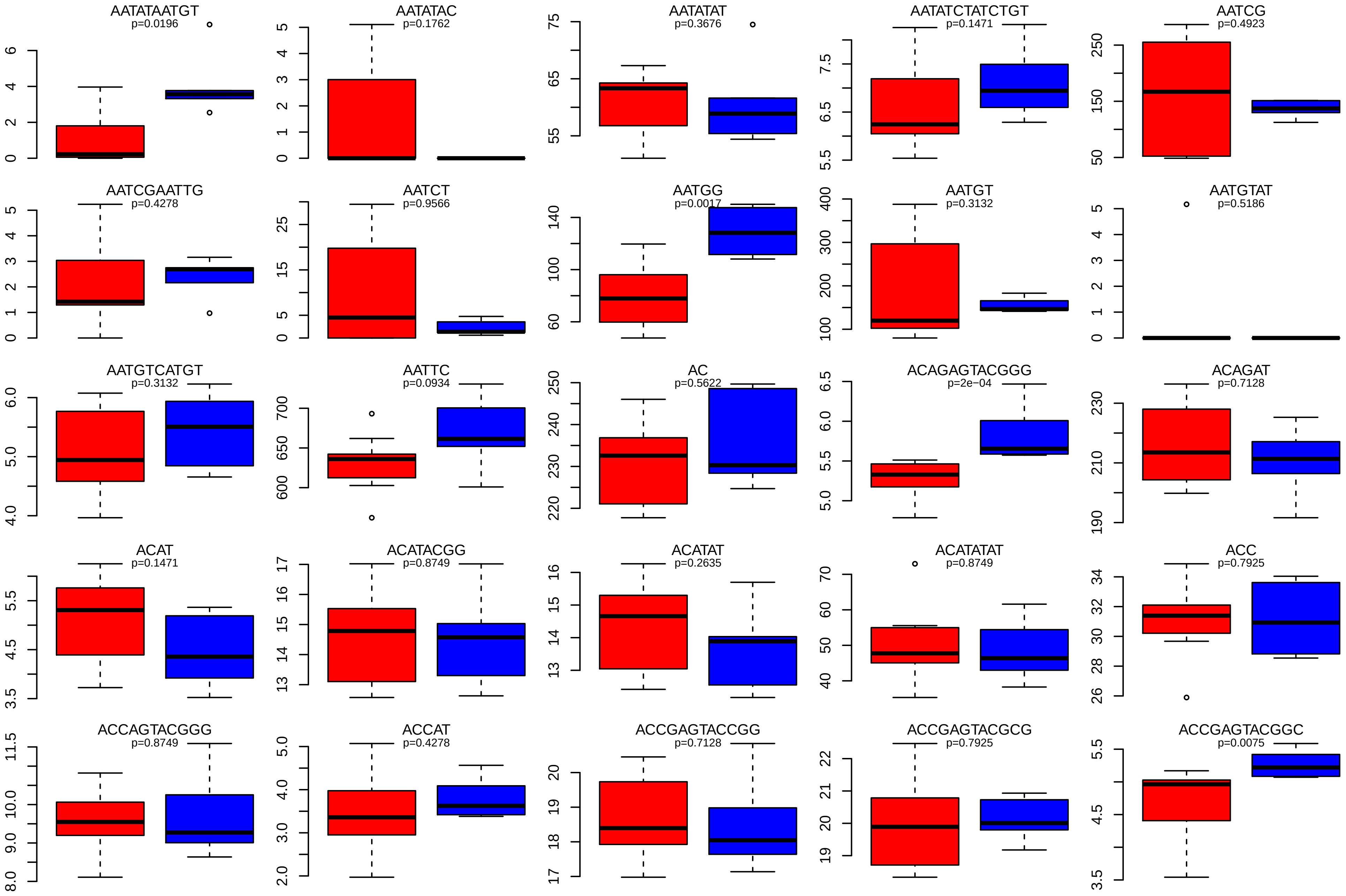

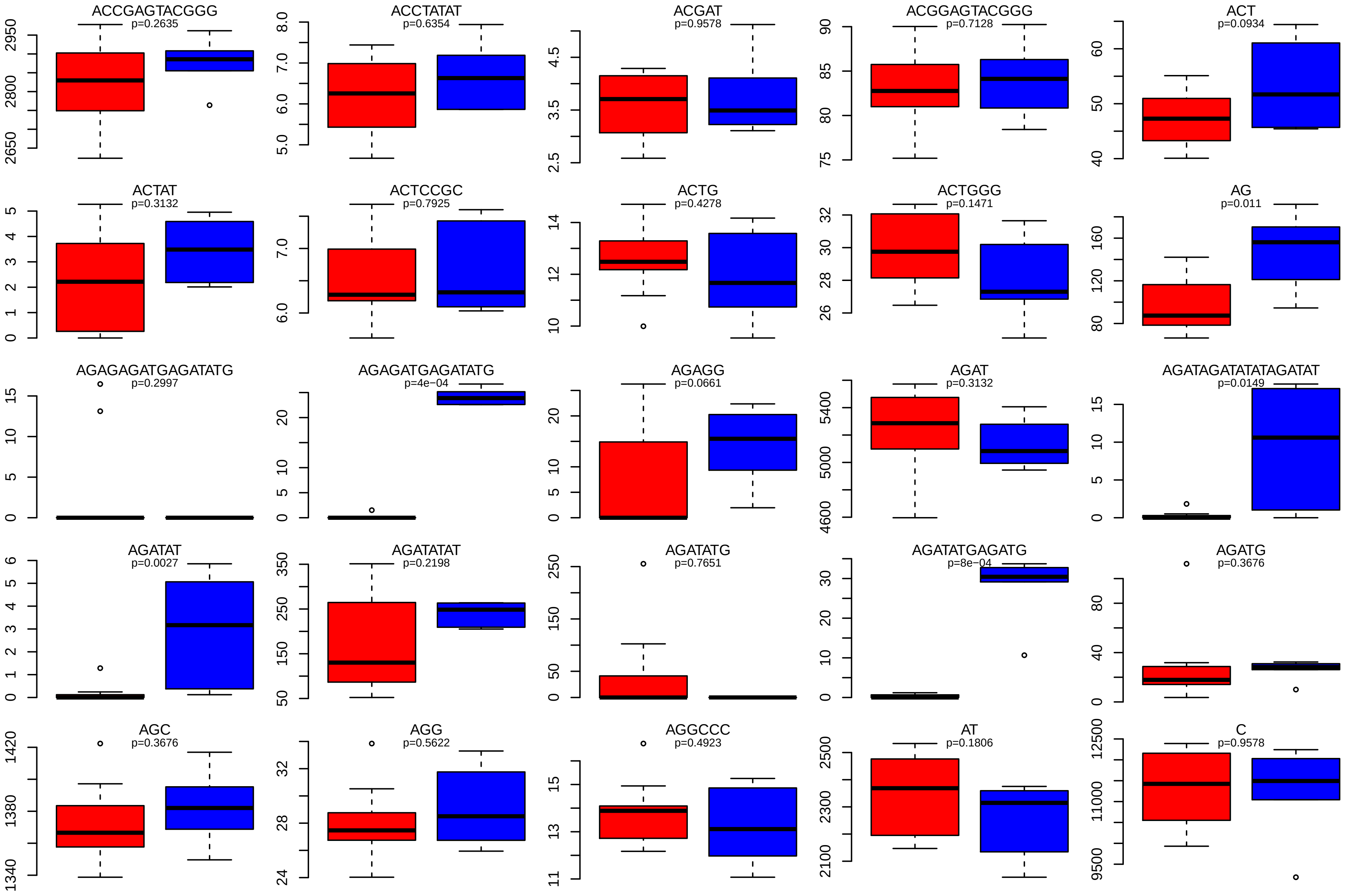

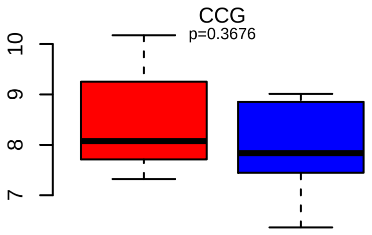

### Supplemental Figure 4

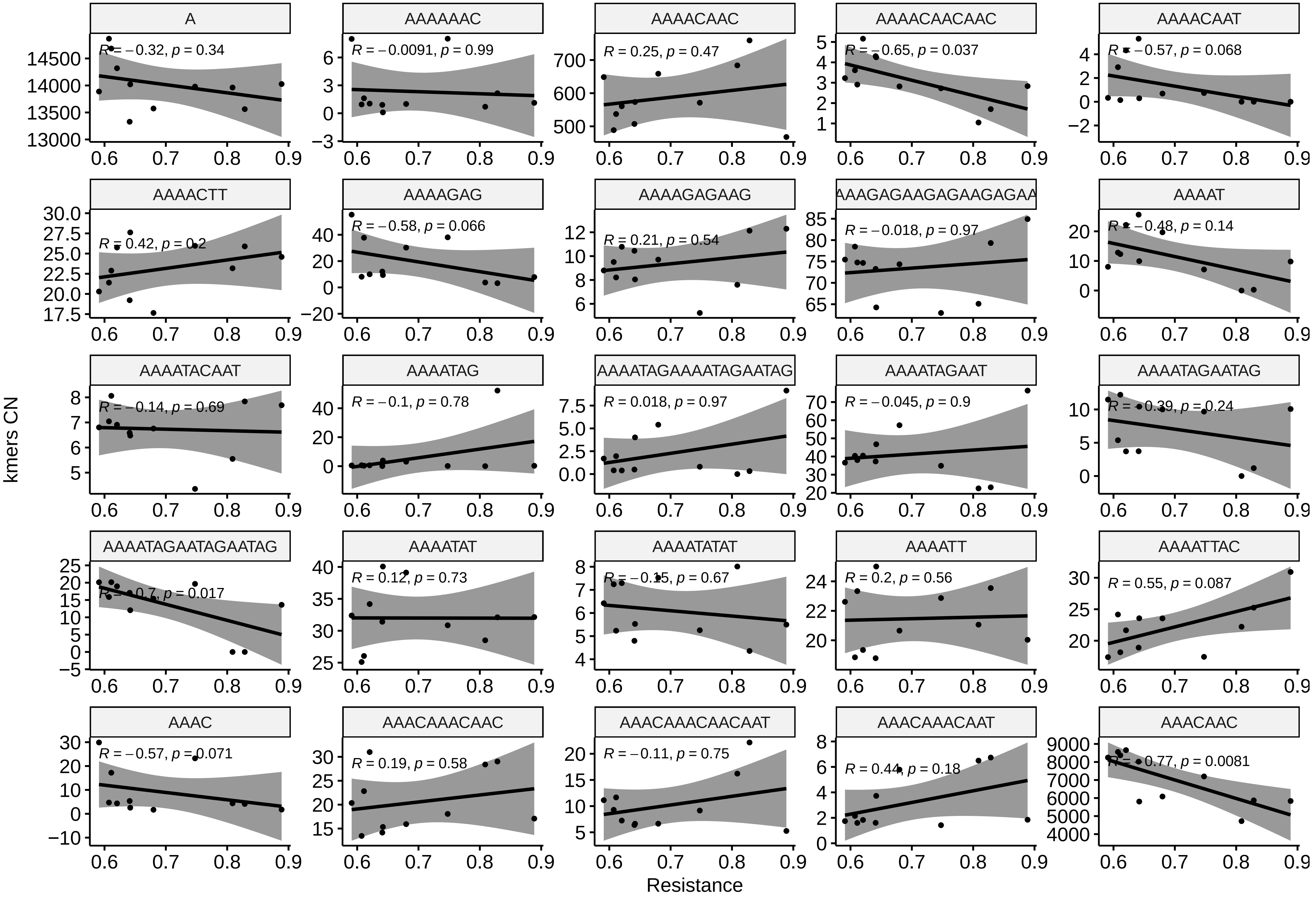

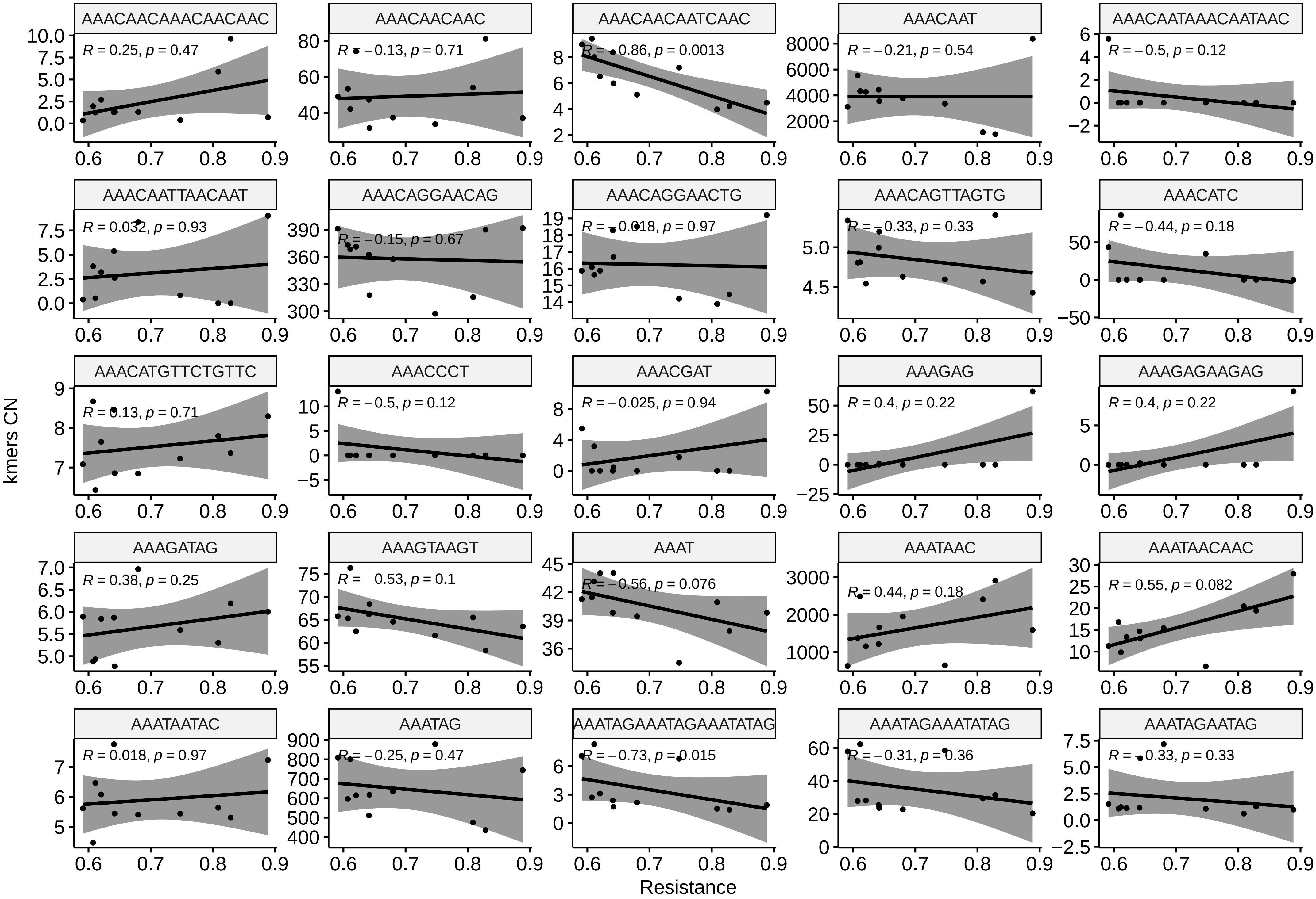

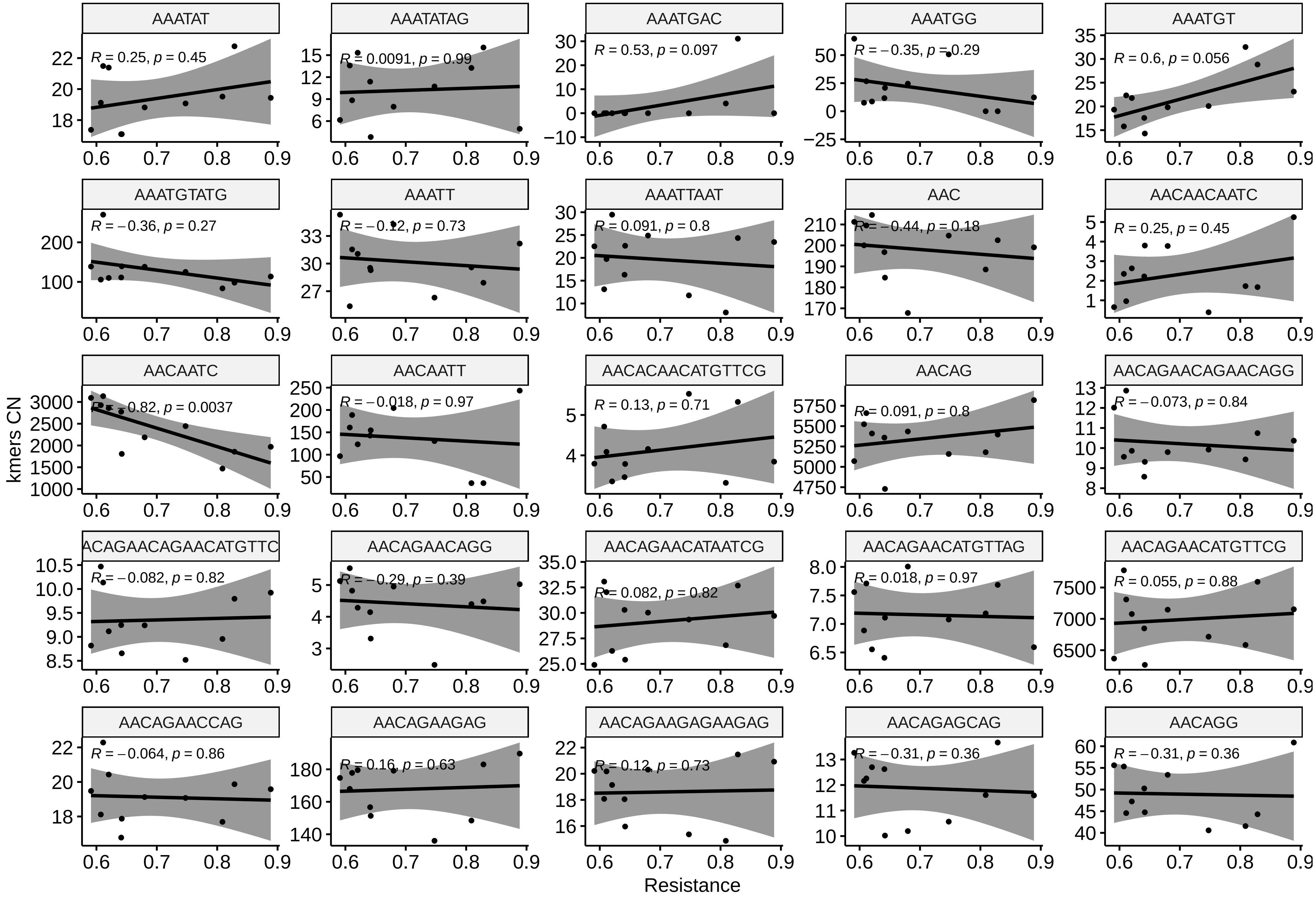

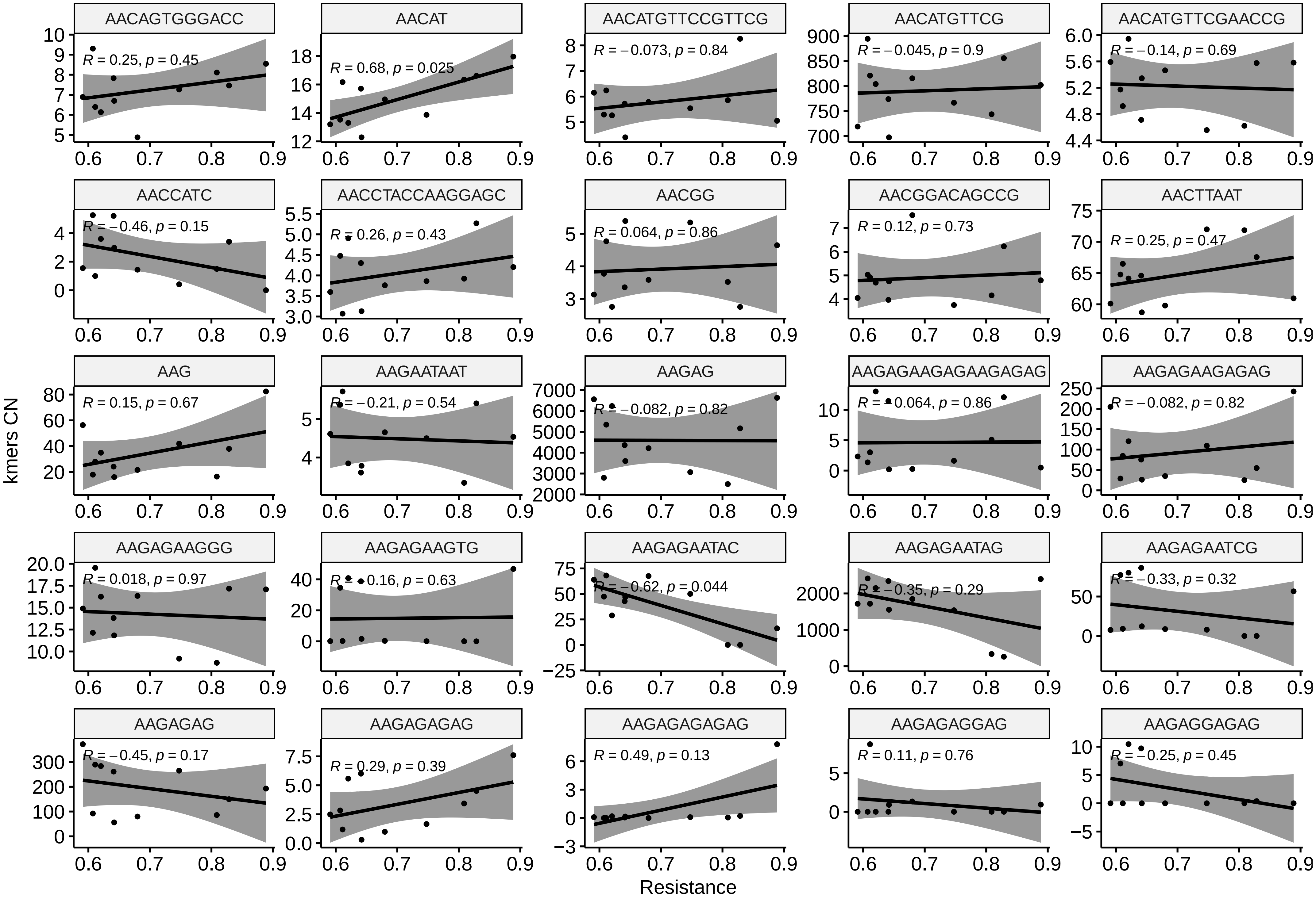

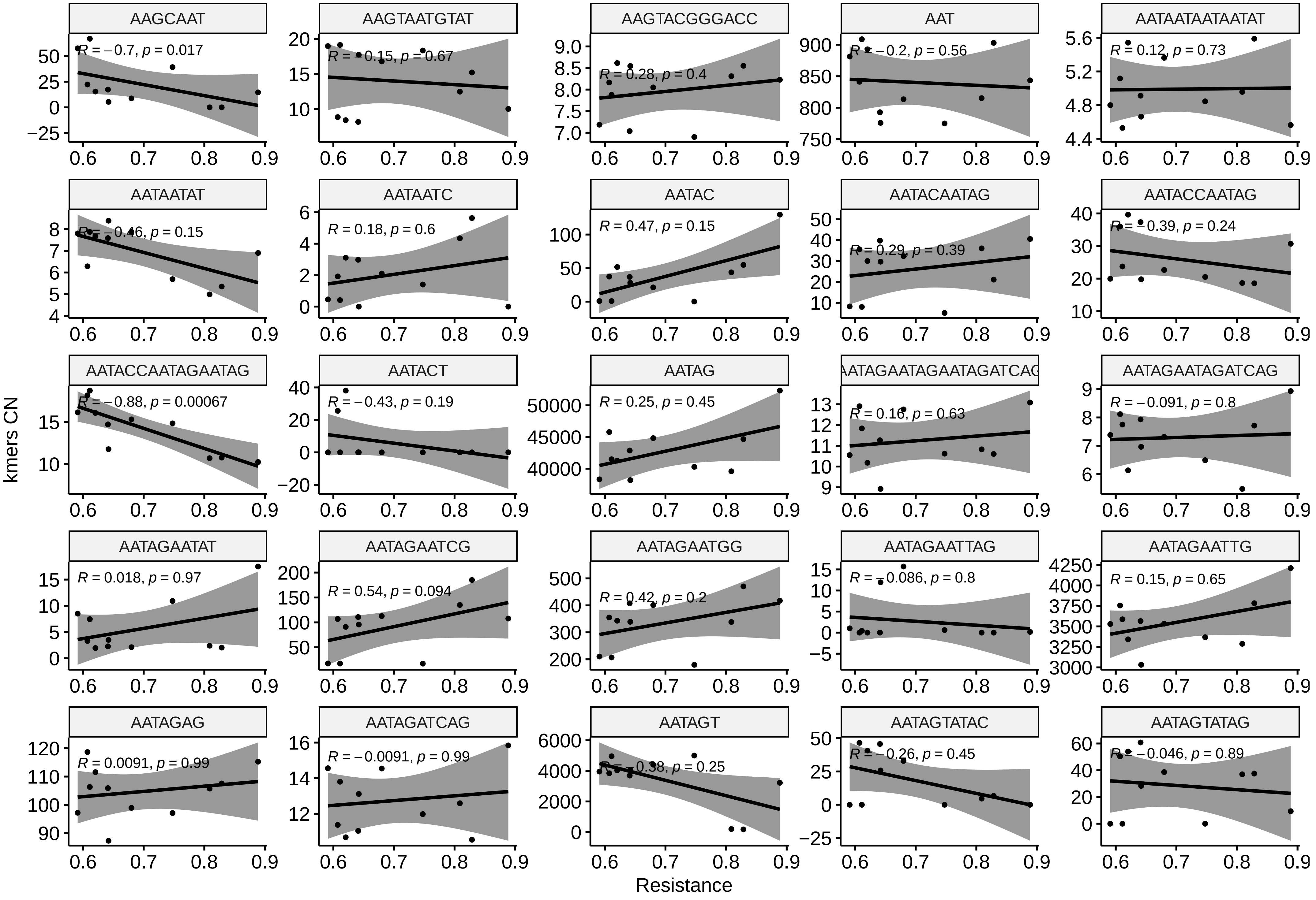

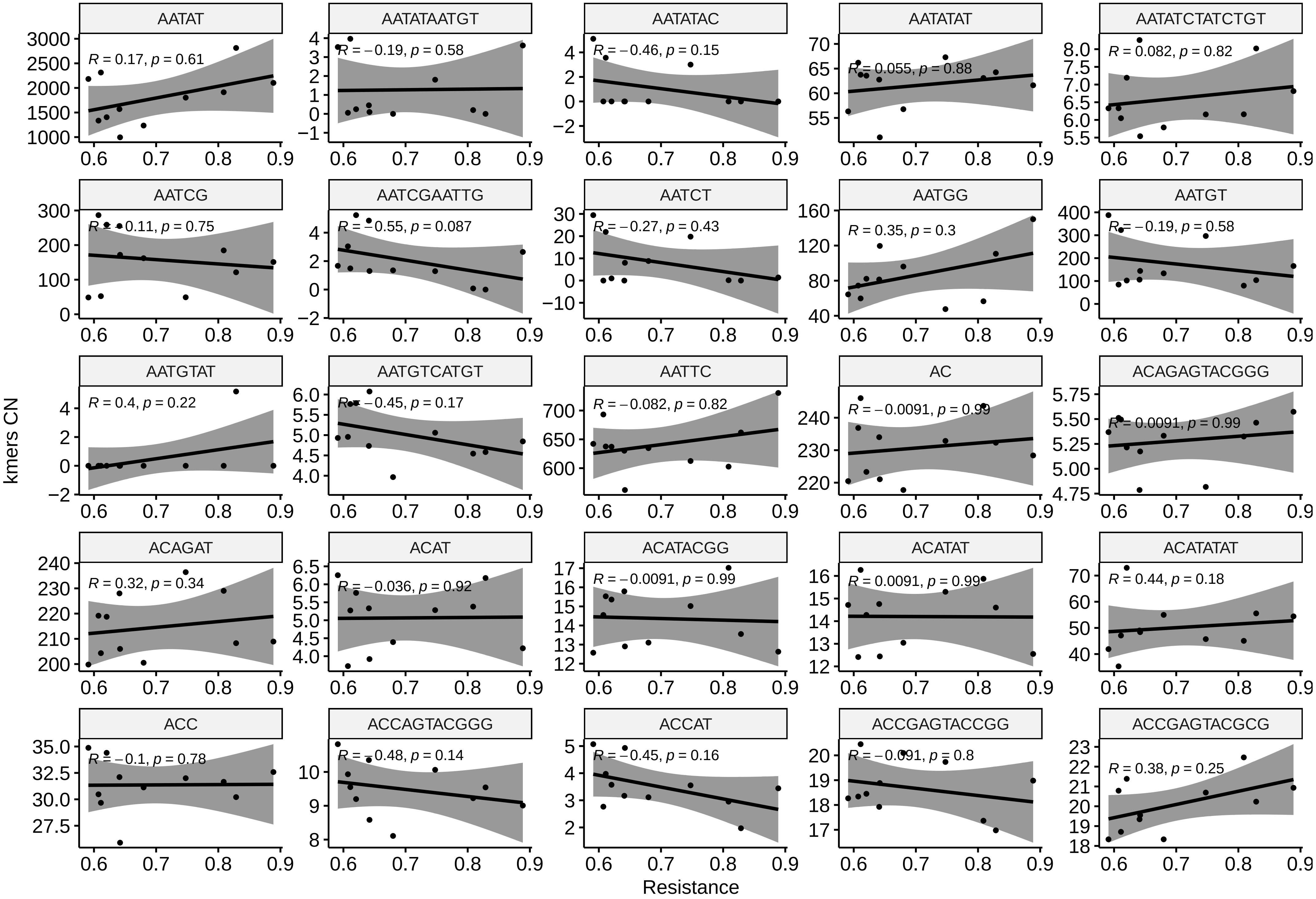

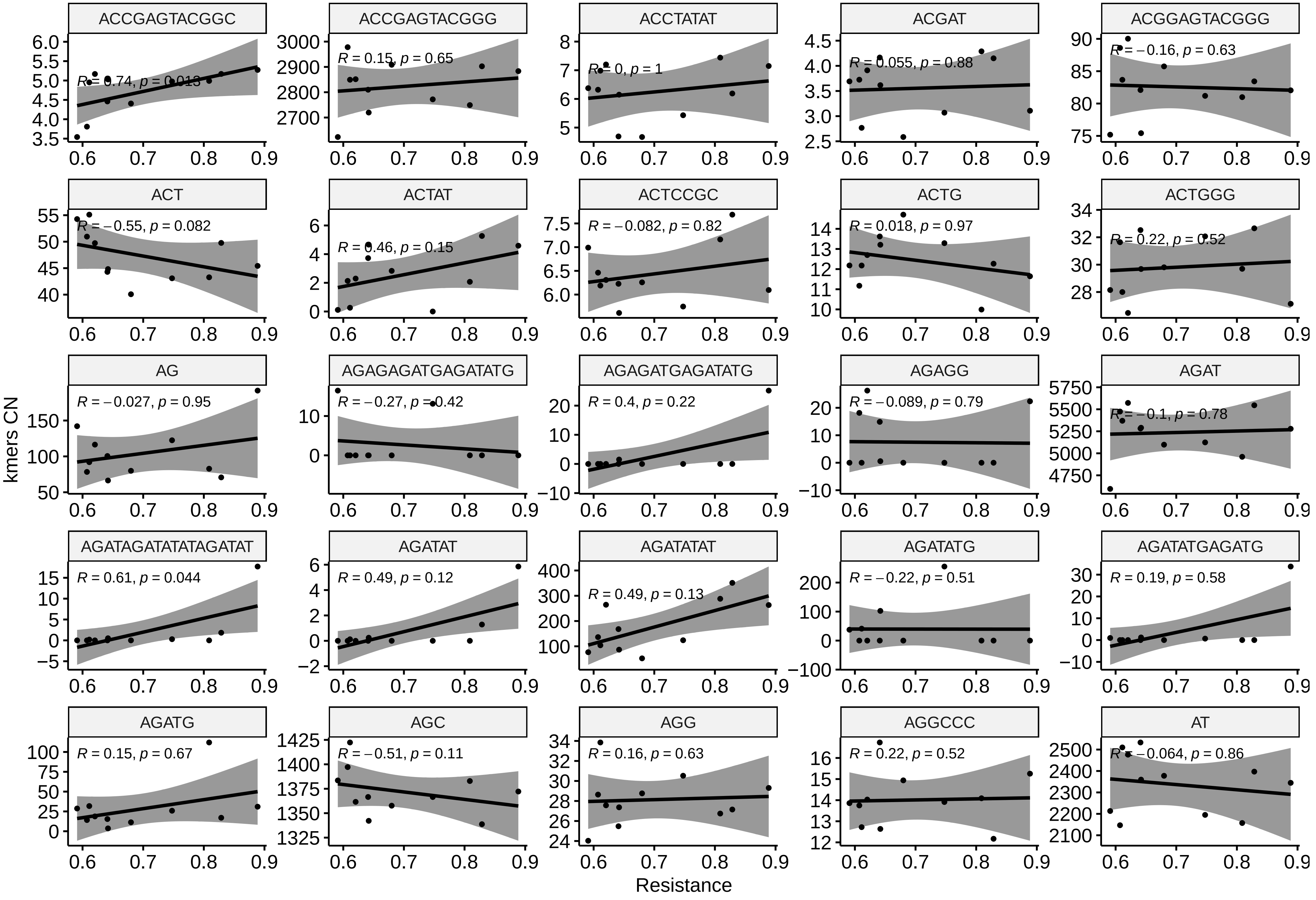

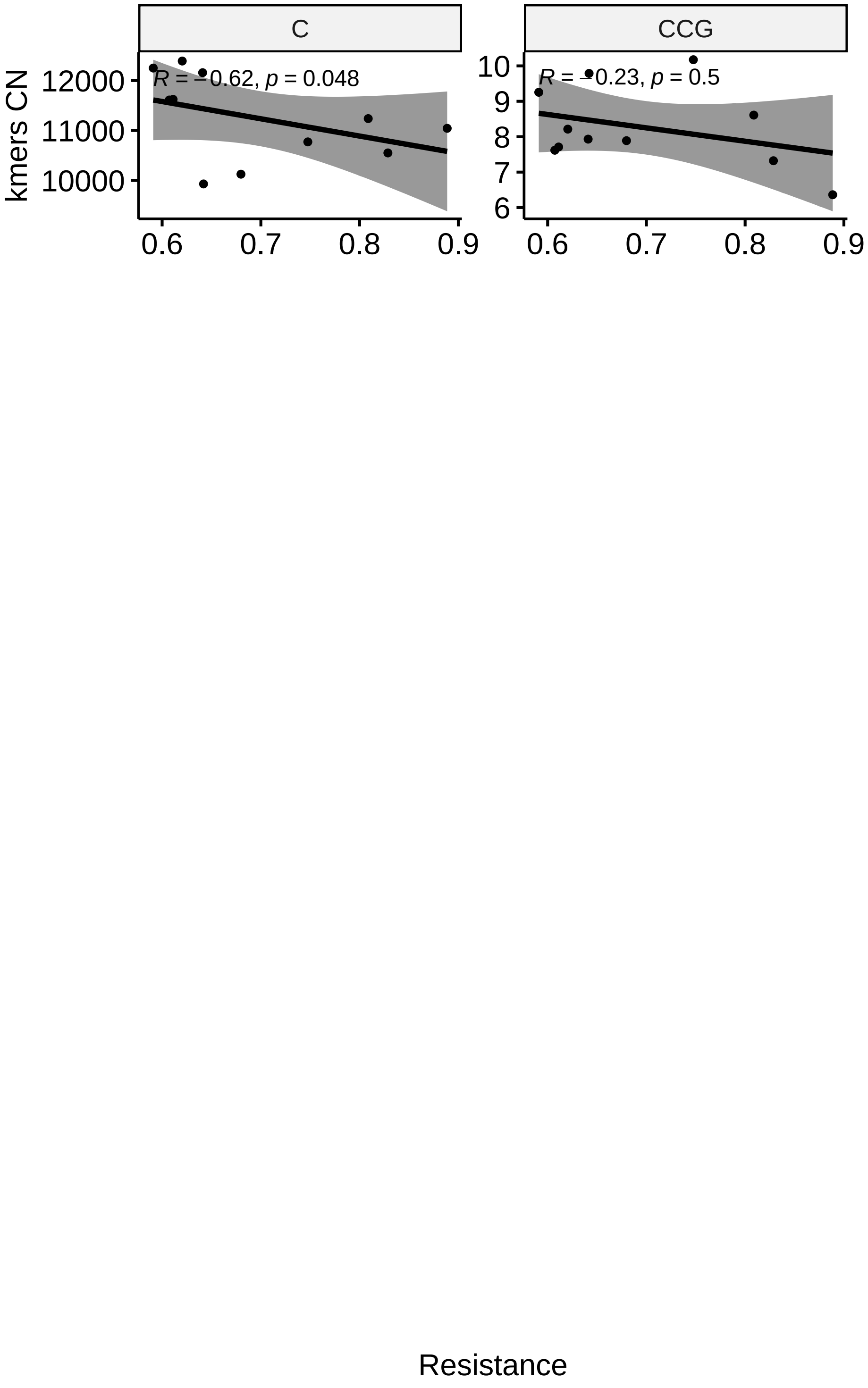

### Supplemental Figure 5

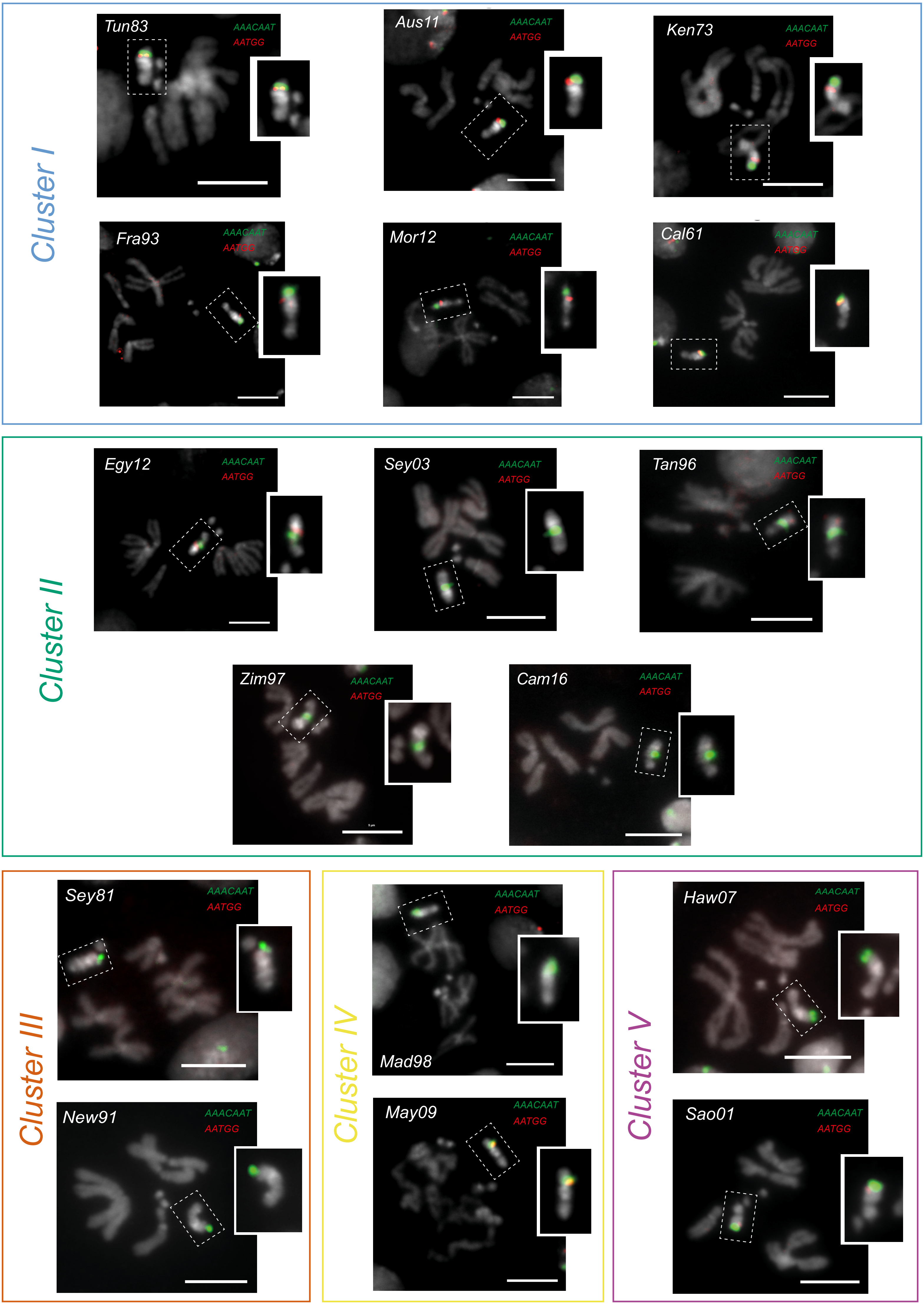

### Supplemental Figure 6

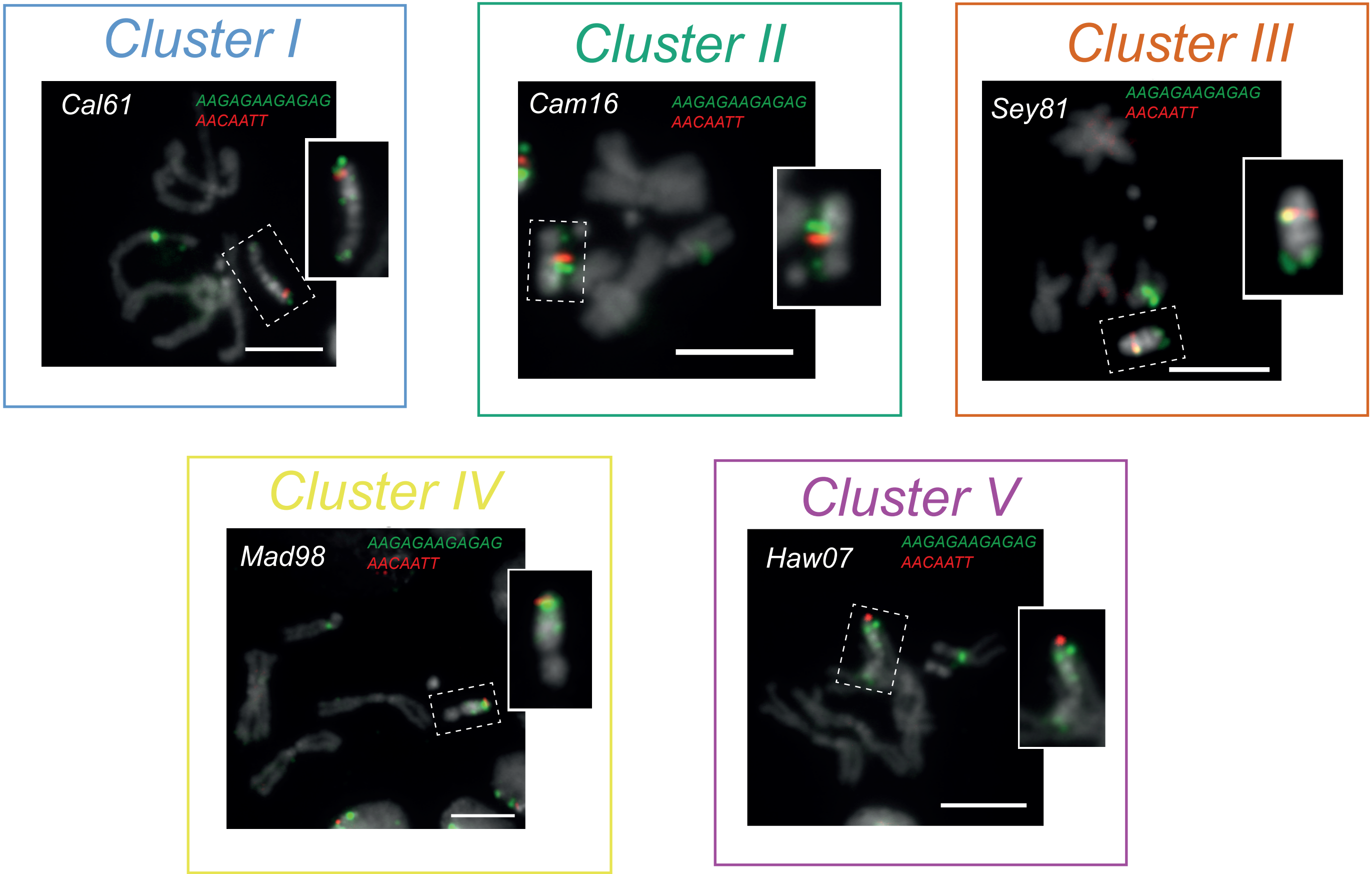

### Supplemental Figure 7

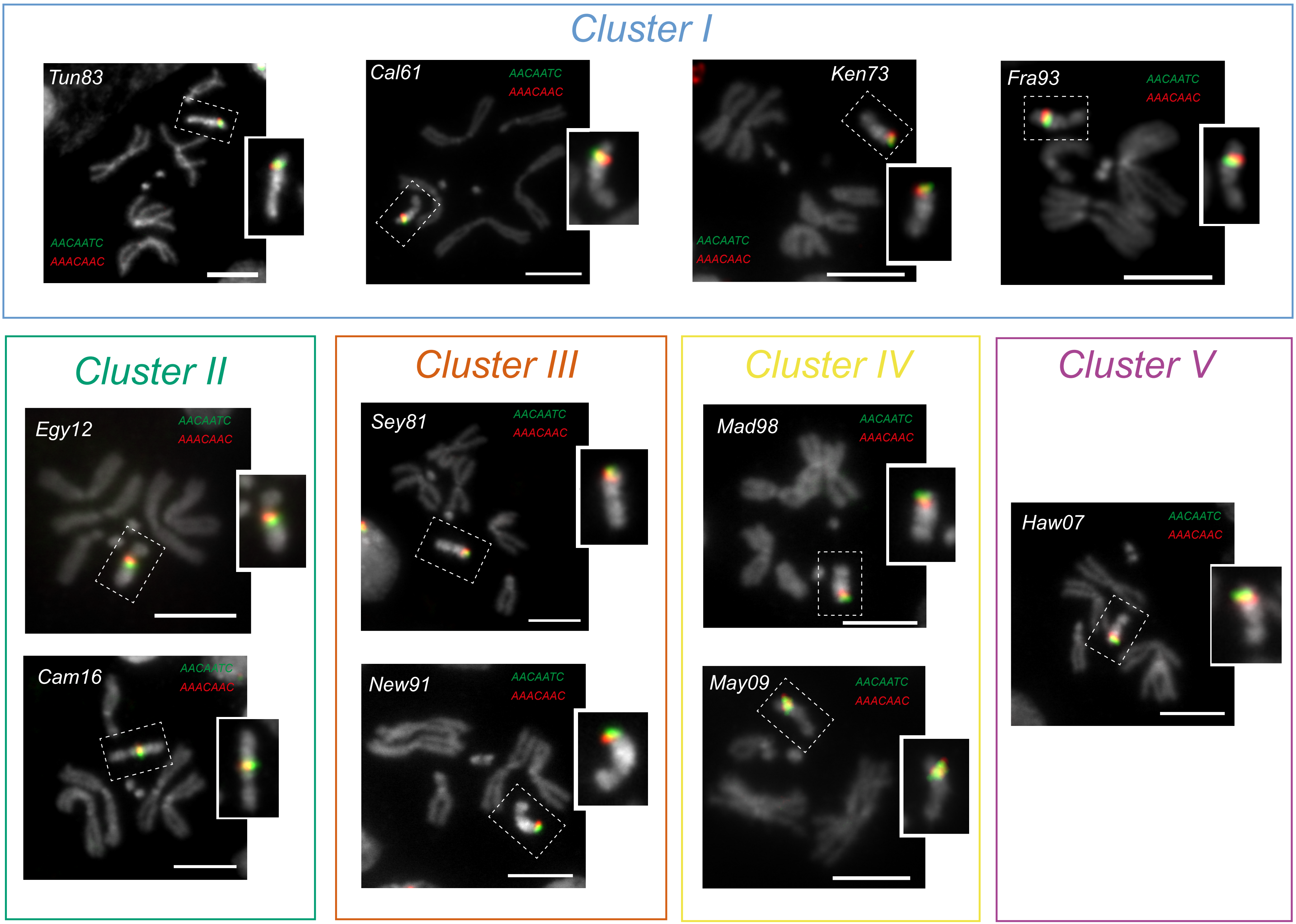

### Supplemental Figure 8

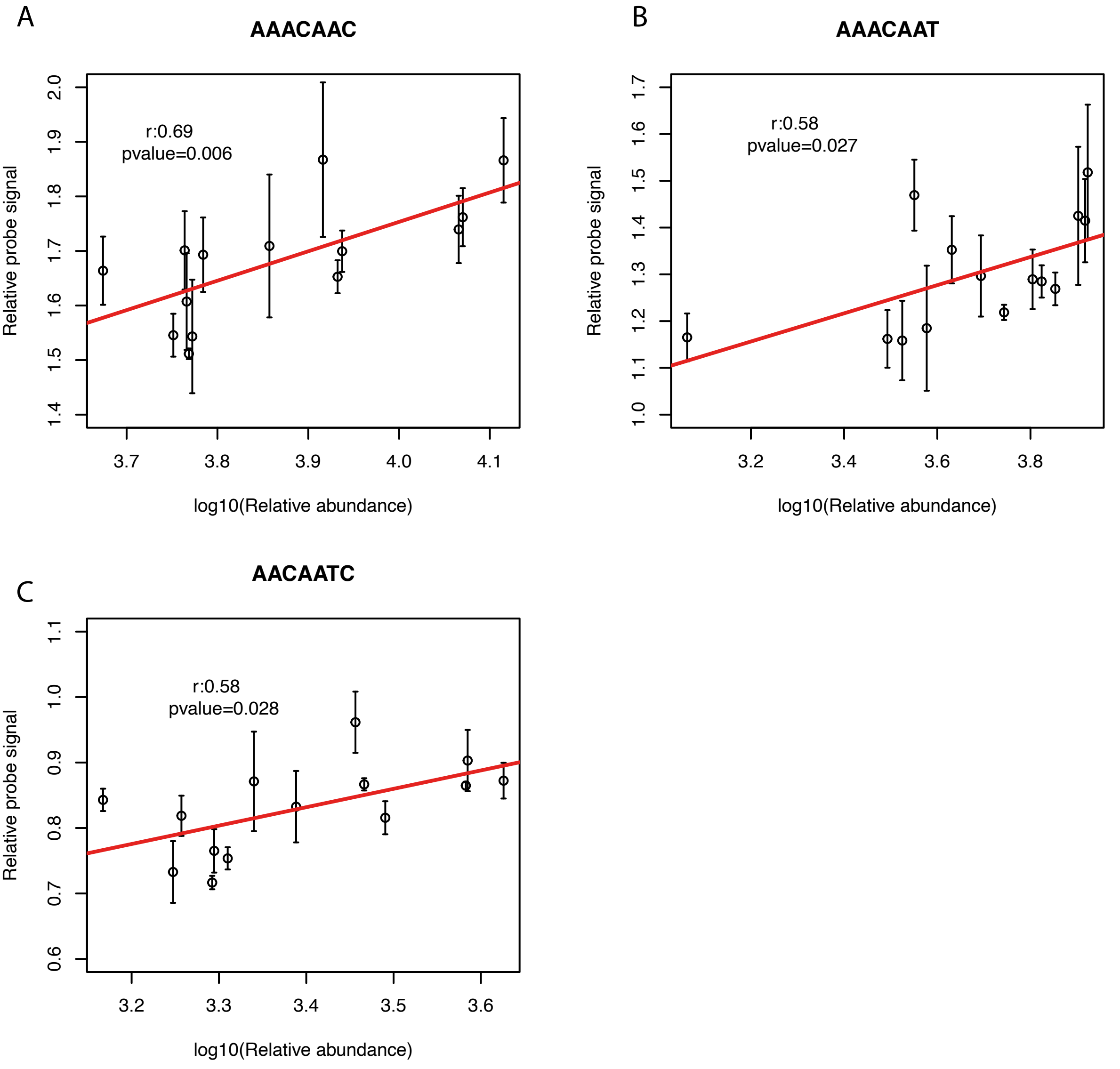
